# A molecular switch in NAC prevents mitochondrial protein mistargeting by SRP

**DOI:** 10.1101/2025.07.29.667405

**Authors:** Emir Maldosevic, Radoslaw Gora, Liangguang Leo Lin, Linyao Elina Zhou, Zexin Jason Li, Yelena Peskova, Ling Qi, Shu-ou Shan, Ahmad Jomaa

**Affiliations:** Department of Molecular Physiology and Biological Physics, University of Virginia, Charlottesville, VA 22903, USA; Department of Biochemistry and Molecular Genetics, University of Virginia, Charlottesville, VA 22903, USA; Medical Scientist Training Program, University of Virginia, School of Medicine, Charlottesville, VA 22903, USA; Division of Chemistry and Chemical Engineering, Caltech, Pasadena, CA 91125, USA

## Abstract

The nascent polypeptide-associated complex (NAC) co-translationally screens all nascent proteins and regulates their access to the signal recognition particle (SRP) to ensure the fidelity of protein targeting to the endoplasmic reticulum (ER). However, the mechanism by which NAC prevents the mistargeting of nascent mitochondrial proteins remains unclear. Here, we identified a molecular switch in NAC that allows its central barrel domain to adopt a stabilized conformation on ribosomes exposing a mitochondrial targeting sequence (MTS). Mutations of the MTS on the nascent chain or in the NAC switch region increases NAC barrel dynamics and reduces its binding to the ribosome. This leads to an impaired ability of NAC to prevent mistargeting by SRP and causes ER stress in human cells. Our work reveals how NAC detects nascent mitochondrial proteins early in translation and prevents their promiscuous access to SRP, elucidating the structural basis that underlies this role and providing novel insights into protein targeting fidelity with broader implications for cellular proteostasis.

## Main

During protein synthesis, nascent polypeptides are co-translationally sorted to distinct cellular destinations, a process critical for maintaining protein homeostasis^1,2^. This process begins on the ribosome, where the nascent polypeptide-associated complex (NAC) binds and screens all emerging nascent chains to regulate their localization and processing^3,4^. The majority of mitochondrial proteins are nuclear encoded and synthesized in the cytosol^5–7^ where they must be properly screened by NAC to ensure accurate targeting. As part of this screening, nascent chain access to targeting factors, such as the signal recognition particle (SRP) that delivers translating ribosomes to the endoplasmic reticulum (ER), must be tightly controlled by NAC to prevent the mistargeting of mitochondrial proteins^8^. Despite its critical role in nascent chain sorting, whether and how NAC detects nascent mitochondrial precursor proteins and distinguishes them from ER-targeted proteins remain unclear.

NAC is an essential and conserved ribosome associated complex in eukaryotes^9–11^ that is expressed at equimolar levels to ribosomes^12^. It is composed of the NACα and NACβ subunits^13^, which dimerize to form a central β-barrel domain^14,15^. NACβ anchors the complex to the ribosome with nanomolar affinity^16^ using a basic N-terminal tail that interacts with both ribosomal proteins and ribosomal RNA (rRNA)^17–19^. Previously, AlphaFold models fitted into lower resolution cryo-EM densities (∼6-8 Å) suggested that two amphipathic helices (βh2 and αh1) in the NAC barrel domain lock together to form a clasp coordinated by a series of hydrophobic residues^19^ which mediate binding of the barrel domain near the polypeptide exit tunnel^17,18^. This positions the barrel domain to potentially interact with nascent chains emerging from the exit tunnel^20,21^ and coordinate their co-translational processing^22–24^ or targeting^19,25,26^.

One of the best-established functions of NAC is to prevent the mistargeting of nascent proteins to the ER^8^ by regulating the interaction, conformation, and activity of SRP^16,19,25,26^. SRP contains an M-domain, which recognizes an N-terminal hydrophobic ER targeting signal on nascent proteins emerging from the ribosome, and a GTPase NG-domain that dimerizes with the SRP receptor (SR) at the ER membrane^16,25,27^. Engagement of the M-domain with the signal sequence activates SRP for rapid assembly with SR to turn on ER targeting^28,29^ and prevents ER proteins from being mistargeted to mitochondria^30^. However, the specificity of ribosome binding and activation of SRP and SR is limited^13,16^. NAC uses dual mechanisms to regulate ER targeting. A flexibly tethered C-terminal UBA domain in NACα recruits SRP to translating ribosomes^19^, whereas the NAC barrel domain prevents the promiscuous activation of SRP by blocking the access of the SRP M-domain to the ribosome exit tunnel^8,25,26^. When an ER signal sequence emerges from the ribosome, the NAC barrel becomes destabilized, allowing SRP to displace it from the ribosome tunnel exit so that the nascent chain can be accurately targeted to the ER^19^. Unlike ER proteins, mitochondrial proteins do not destabilize NAC binding to the ribosome^19^. Thus, the specificity of nascent chain sorting by NAC is crucial to prevent mitochondrial protein mistargeting to the ER^8^.

A significant subset of mitochondrial proteins contain an N-terminal mitochondrial targeting sequence (MTS) that allows for their proper subcellular localization^31,32^. An MTS is a bipartite sequence motif characterized by clusters of positively charged residues on the polar face of a short amphipathic helix followed by a disordered sequence of 20-90 amino acids. The MTS, in particular the N-terminal basic amphiphilic helix, is recognized by receptors and the translocase at the outer mitochondrial membrane (TOM) and is often necessary and sufficient to direct nascent mitochondrial proteins for targeting and import^33–35^. However, whether NAC directly senses the MTS during translation prior to targeting and how NAC prevents promiscuous interactions of the MTS with SRP on the ribosome remains poorly understood.

To address these questions, we determined the cryo-EM structure of human NAC bound to ribosomes translating a mitochondrial nascent chain with an N-terminal MTS. The structure reveals that the NAC barrel domain shifts to adopt a stabilized conformation near the polypeptide exit tunnel mediated by a new set of ribosomal contacts that are distinct from previous observations of NAC bound to ribosomes translating an ER signal sequence or a cytosolic protein^19,22,23^. We identified a molecular switch in NACβ that allows NAC to alternate between different conformations. Single-molecule studies and biochemical assays indicate that mutations of the MTS disrupt the anchoring of the NAC barrel domain at the ribosome exit site, leading to the promiscuous activation of SRP on ribosomes exposing a nascent mitochondrial protein. Finally, we show that mutations in the NAC switch region led to ER stress phenotypes that are associated with protein mistargeting in human cells. Our work uncovers a previously unappreciated ability of NAC to detect an MTS early during translation and the molecular mechanism employed by structural elements within NAC to prevent the mistargeting of nascent mitochondrial proteins.

## Results

### The NAC barrel adopts a distinct conformation on ribosomes translating an MTS

Previous studies established that NAC mediates proper protein sorting by preventing SRP access to ribosomes that do not carry signal sequences^8,19,25^. To first confirm if NAC deletion leads to mitochondrial protein mislocalization in human cells, NACβ knockout (KO) HEK293T cells were generated using CRISPR-Cas9 (Extended Data Fig. 1A). Immunofluorescence staining was conducted to assess the localization of Oxa1L and Hsp60, which are targeted to the inner mitochondrial membrane and mitochondrial matrix, respectively. In wild-type (WT) cells, both Hsp60 and Oxa1L co-localized with the mitochondrial marker Tom20 (Extended Data Fig. 1B-C). In contrast, NACβ KO cells showed significant mislocalization of both proteins, with fluorescence signals accumulating outside mitochondria, indicating impaired mitochondrial protein sorting in the absence of NACβ.

Next, we asked how NAC engages with ribosomes carrying an MTS. Model MTS-containing nascent chains derived from Oxa1L and Hsp60 were generated by stalling ribosomes during translation with the XBP1 arrest peptide^36^ or on mRNA lacking a stop codon^37^, respectively. Purified mammalian ribosome-nascent chain complexes (RNCs) exposing the N-terminal MTS (RNC_MTS_) of Oxa1L (74 residues) or Hsp60 (62 residues) were incubated with human NAC. As a control, an MTS deletion mutant of Oxa1L was also purified (Extended Data Fig. 2). Single particle cryo-EM analysis was used to visualize the interactions between NAC and ribosomes translating mitochondrial nascent chains (Extended Data Table 1).

Initial 2D and 3D classifications yielded heterogenous 80S particles with different translation states. Further iterative 3D variability analyses were conducted using focused masks to sort for ribosomes translating mitochondrial nascent chains, which resulted in a final set of particles containing density for P-site tRNA, nascent chain, and NAC (Extended Data Fig. 3A-B, Extended Data Fig. 4A-B). The cryo-EM structures of both Oxa1L- and Hsp60-RNC_MTS_ resolved NAC bound next to the polypeptide exit tunnel. The overall resolution of the two maps was ∼2.6 and ∼3.0 Å. NAC was resolved between 3-6 Å (Extended Data Fig. 3C-D, Extended Data Fig. 4C-D) in a similar conformation on the ribosome for both MTS constructs (Extended Data Fig. 5A-C). Since the NAC barrel domain was better resolved in the Oxa1L-RNC_MTS_ map, this map was used for further interpretation of the structure.

The NAC barrel was docked near the polypeptide exit tunnel (red asterisk, Fig. 1A), consistent with its role in protecting and scanning emerging nascent chains^8,21^. Cryo-EM density corresponding to the nascent chain was present in the polypeptide exit tunnel, but the nascent chain including the MTS was not resolved outside the ribosome due to flexibility (Figure 1A, Extended Data Fig. 6A-B). In the current structure, NAC was stabilized by a series of contacts between NACβ and the ribosome (Fig. 1B-E) that were not previously observed in the structure of NAC bound to ribosomes translating an ER signal sequence (RNC_SS_)^19^.

**Figure 1.**
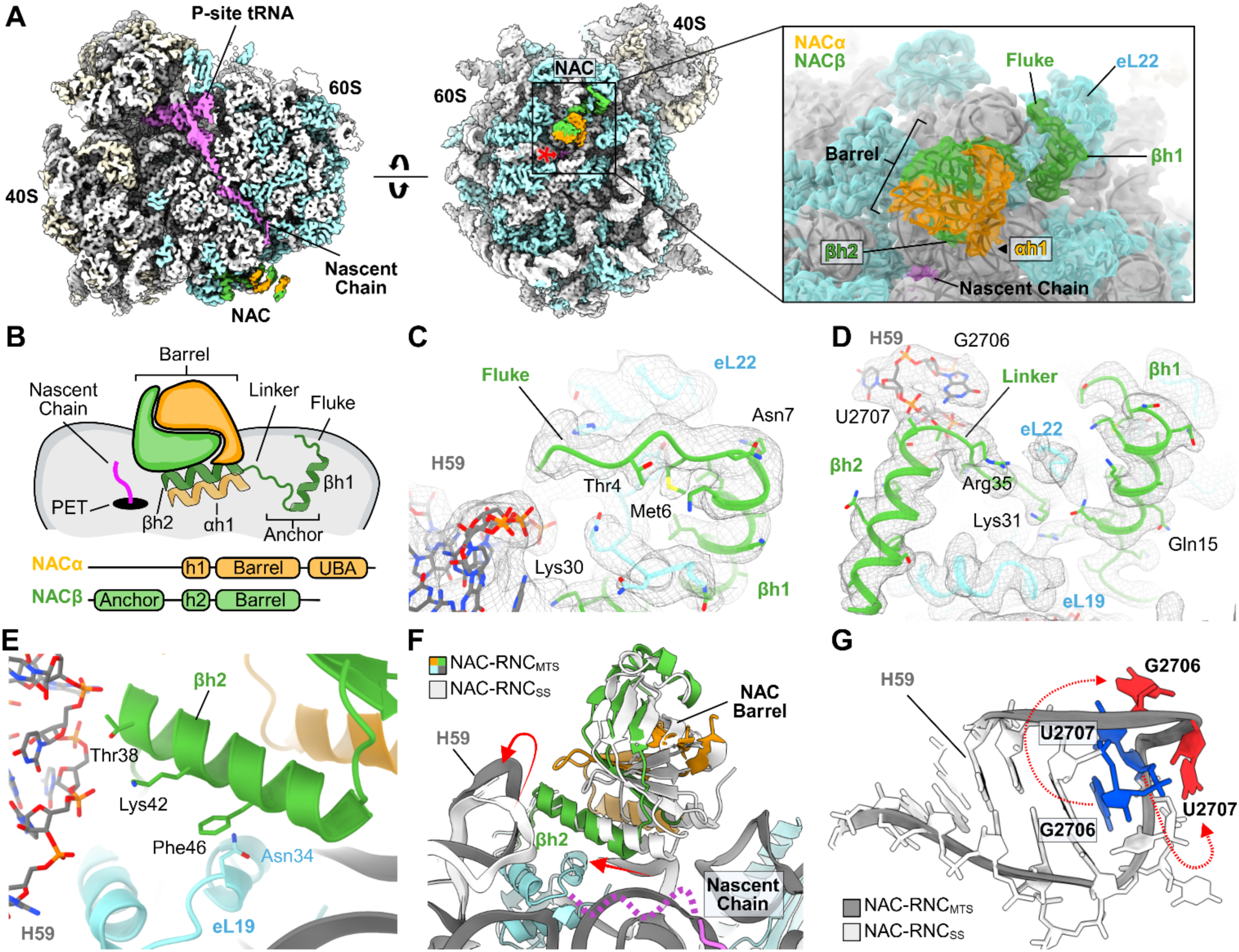
Structure of NAC in complex with a ribosome translating a mitochondrial nascent chain. **A)** Cryo-EM structure of the Oxa1L NAC-RNC_MTS_ complex. The inset shows a closeup of NAC docked near the polypeptide exit tunnel (PET-red asterisk). Large ribosomal proteins are colored in cyan; small ribosomal proteins are colored in beige; NACβ is colored in green, and NACα is colored in orange; P-site tRNA and the nascent chain are colored pink; rRNA is colored in grey. **B)** Schematic of NAC bound near the PET. **C)** Interaction of the N-terminal fluke of NACβ with eL22 and H59, with the cryo-EM map shown as mesh. **D)** Interaction of NAC βh1, βh2, and the intervening linker with rRNA H59 and eL19. The cryo-EM map is shown as mesh. **E)** Side view of NAC βh2 contacts with eL19 and H59. **F)** Comparison of the NAC-RNC_MTS_ model to the previously published model of ribosomes translating an ER signal sequence (NAC-RNC_SS_, PDB 7QWR). Red arrows indicate the conformational change observed for the NAC barrel and H59. **G)** Rearrangement of H59 between the NAC-RNC_MTS_ and NAC-RNC_SS_ structure. Red arrows depict the conformational change of the bases G2706 and U2707 in the RNC_MTS_ model (red), compared to the RNC_SS_ model (blue).

NACβ contains a conserved N-terminal RRKKK motif, which is essential for high-affinity ribosome binding of the complex^17^. This ribosome ‘anchor’ of NAC was previously observed to intercalate around the C-terminal tail of eL22, forming extensive interactions between the ribosomal RNA (rRNA) and the ribosomal protein eL19^19^. In addition to the previously observed ribosome contacts, we resolved an additional insertion of the first 4 N-terminal residues of NACβ wedged between rRNA helix 59 (H59) and eL22, which forms the fluke of the NAC anchor (Fig. 1C, Extended Data Fig. 7A-B). A combination of polar and stacking interactions mediate the contacts with eL22, consistent with crosslinking data^21^.

The newly resolved contacts are established following a large conformational shift of the NAC barrel towards H59 (colored grey vs. white, Fig. 1F). In particular, the base of the barrel that directly contacts the ribosome moves by 10 Å away from the polypeptide exit tunnel. To accommodate this shift, H59 undergoes a conformational change where it lifts away from the ribosome by 9 Å (red arrows, Fig. 1F). In this state, the rRNA bases G2706 and U2707 are flipped out following rearrangement of the rRNA (red arrows, Fig. 1G).

Two predicted amphipathic helices in the NAC barrel domain (βh2 and αh1) and part of the linker on NACβ that connects the barrel domain to the anchor (Fig. 1B) were resolved at side-chain resolution for the first time and built de novo (Fig. 1D and Extended Data Fig. 8A-C). The two amphipathic helices form a clasp containing a hydrophobic core (Extended Data Fig. 8A) and mediate the interactions (described below) that stabilize the NAC barrel on the ribosome. These interactions are distinct from those in the previous structures of NAC bound to ribosomes translating an ER signal sequence, in which both the hydrophobic clasp and contacts with the ribosome were destabilized by the emergence of a signal sequence and thus were not well resolved^19^.

### A molecular switch regulates NAC binding to the ribosome

Several residues in NAC βh2 and the preceding linker, conserved among higher eukaryotic organisms, were identified as mediating interactions with the ribosome (Fig. 2A). In the NACβ linker region, His34 inserts below H59 to lock NAC in a stable conformation on the ribosome (Fig. 2A), while the side chain of Arg35 interacts with the flexible C-tail of eL22 (Fig. 2B and Extended Data Fig. 8A). Furthermore, Lys42 is within range for electrostatic interactions with the phosphate backbone of rRNA H59, while Phe46 stacks with Asn34 on eL19 (Fig. 1E, Extended Data Fig 8C). These contacts, which were not observed in the NAC-RNC structure containing ER-destined nascent chains, shifted the NAC barrel position at the polypeptide exit tunnel compared to the previous structure (Fig. 1F, Fig. 2B).

**Figure 2.**
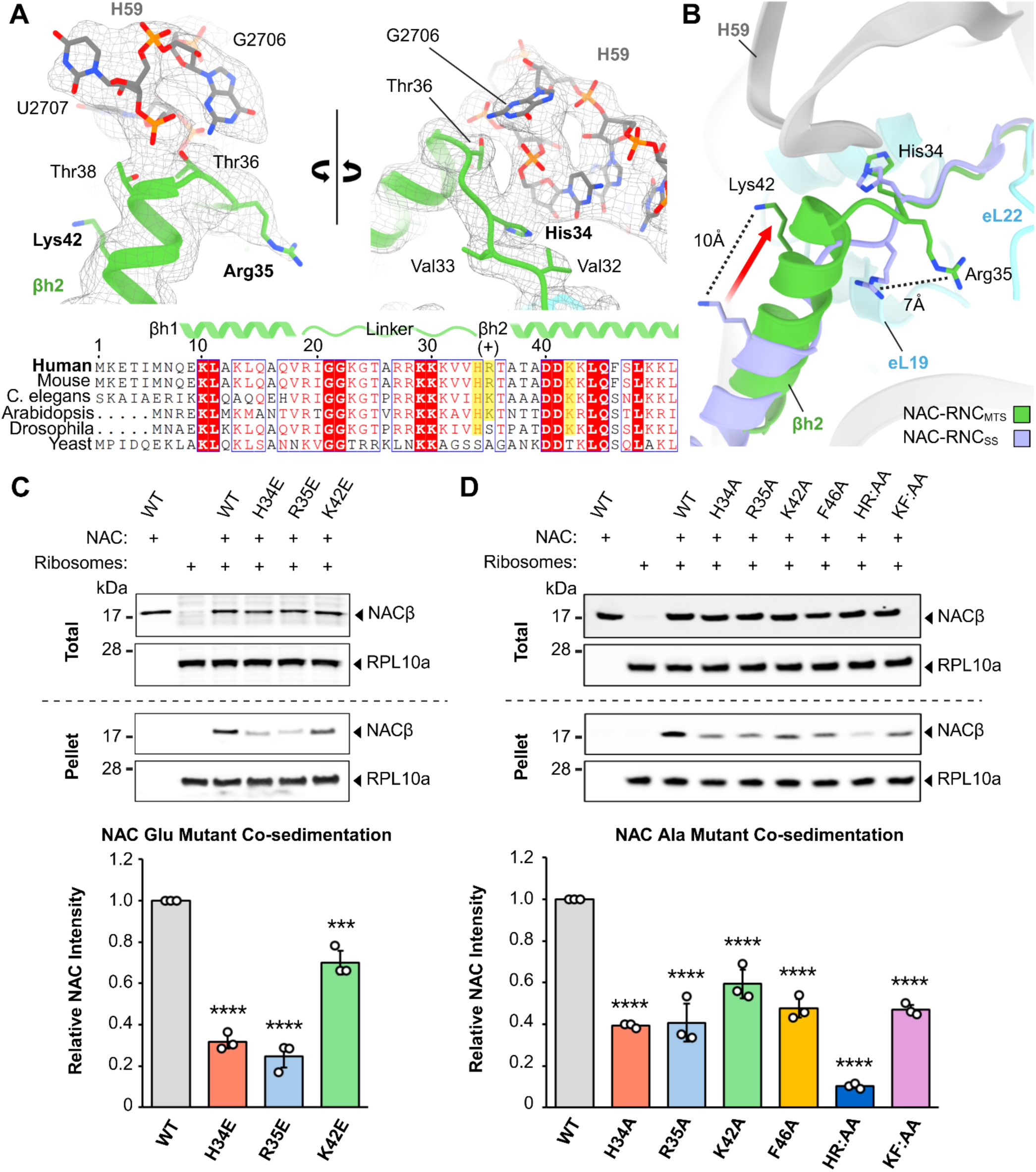
Structural and biochemical analysis identifies a molecular switch in NACβ. **A)** Closeup of the residues in NAC βh2 and the preceding linker that mediate interactions with H59. The cryo-EM map is shown as a mesh. Sequence alignments below show the conservation of the highlighted residues/charge. **B)** Comparison of the conformational change in NACβ between the NAC-RNC_MTS_ structure and the NAC-RNC_SS_ (PDB 7QWR), depicted with an arrow. Dashed lines indicate distance of the movement of Lys42 and Arg35. **C, D)** Representative western blots showing the ribosome association of WT NAC and indicated mutants in the NAC switch region determined by the co-sedimentation assay. RPL10a was used as a loading control. Additional replicates used in the quantifications are shown in Extended Data Fig. 9C-D. Bar graphs below the blots show the quantification of NAC bands in the pellet relative to WT NAC (n=3). Values are plotted as Mean ± SD. Statistics: one-way ANOVA with Dunnett’s multiple comparisons test (***p<0.001, ****p<0.0001).

To investigate the role of these novel contacts in ribosome binding, residues mediating NAC barrel domain interactions with the ribosome (His34, Arg35, Lys42, and Phe46) were mutated to either Glu/E or Ala/A (Extended Data Fig. 9A-B). We tested the ability of the NAC mutants to interact with human ribosomes isolated from HEK293 cells using an *in vitro* co-sedimentation assay. Both single and double mutations in the NACβ linker (His34 and Arg35) and the amphipathic helix βh2 (Lys42 and Phe46) significantly reduced NAC association with ribosomes when compared to wild-type (WT) NAC (Fig. 2C-D, Extended Data Fig. 9C-D).

Together, our structural and biochemical data reveal a molecular switch within NACβ that undergoes a conformational change involving the linker residues (His34 and Arg35) and the amphipathic helix βh2. Upon binding ribosomes displaying an MTS, this switch engages with rRNA helix H59 and stabilizes the positioning of the NAC barrel near the polypeptide exit tunnel.

### The molecular switch alters NAC barrel dynamics on the ribosome

We next directly assessed how contacts of the switch region (Fig. 3A) influence NAC conformations on the ribosome by conducting single-molecule total internal reflection fluorescence microscopy (smTIRFM) studies using Oxa1L RNC_MTS_ immobilized on a glass coverslip surface (Fig. 3B). This approach selectively monitors populations of WT or mutant NAC that are bound to surface-immobilized ribosomes and excludes unbound NAC.

**Figure 3.**
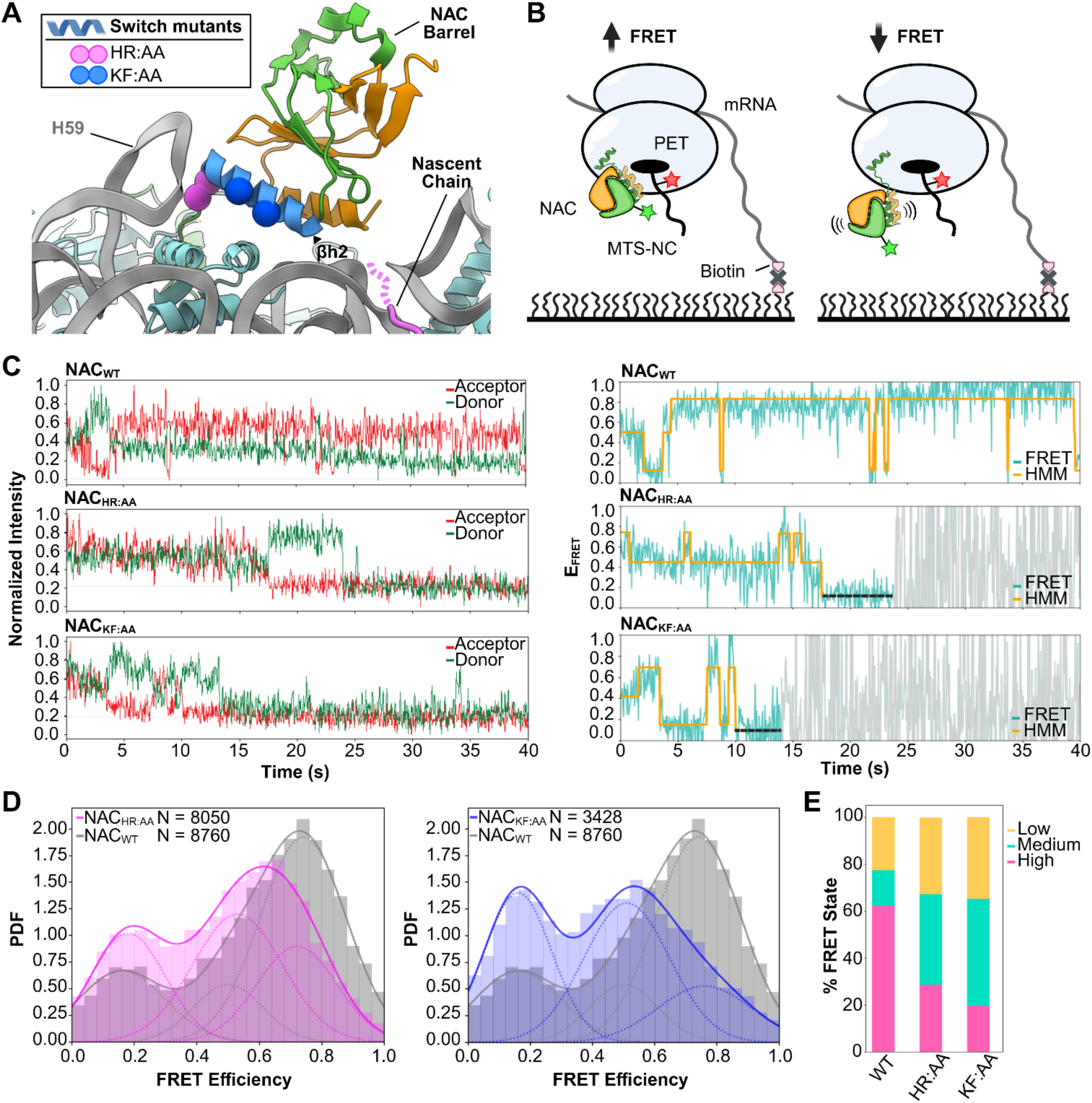
NAC molecular switch mutants impair barrel docking near the ribosome exit tunnel. **A)** NAC-RNC_MTS_ structure, with spheres depicting mutated NAC-ribosome contacts in the switch region used for single molecule experiments. **B)** Schematic representation of the smTIRFM experiment to define NAC barrel dynamics on the ribosome. Oxa1L RNC_MTS_ was immobilized on quartz slides. FRET efficiency between a donor dye (green star) in the NAC barrel and an acceptor dye (red star) on the nascent chain measures NAC barrel docking at the polypeptide exit tunnel (PET). **C)** Representative smFRET traces of NAC interacting with Oxa1L RNC_MTS_. Left panels show the acceptor (red) and donor (green) emission intensity when the donor is excited. Right panels show the corresponding FRET traces (cyan) and Hidden Markov Modeling (HMM) of the data (orange lines) to establish the number of states and their mean FRET values. Dashed black lines indicate acceptor bleaching, while grey indicates the data after donor bleaching; both are excluded from further data processing. **D)** smFRET histograms for wild-type NAC (grey) and mutants NAC_HR:AA_ (left; magenta) and NAC_KF:AA_ (right, blue). PDF, probability density function. ‘N’, number of measurements. The solid lines show the fit of data to the sum of three Gaussian functions with low, medium, and high mean FRET values, and dotted lines show the constituent Gaussian components. **E**) Summary of the distribution of WT and mutant NAC in the different FRET states.

The position of the NAC barrel at the ribosome tunnel exit was measured by Förster resonance energy transfer (FRET) between a donor dye (Cy3B) on the NAC barrel domain and an acceptor dye (Atto647N) incorporated at residue 39 in the Oxa1L nascent chain. This residue is 35 amino acids from the C-terminus of the nascent chain and places the dye at the polypeptide exit tunnel (Fig. 3B). Hidden Markov modeling (HMM) of the fluorescence time traces showed dynamic transitions of the NAC barrel between three states, with low, medium, and high FRET efficiency, on the second timescale (Fig. 3C). The majority of WT NAC displayed long-lived high FRET traces with brief transitions to medium and low FRET states (Fig. 3C), indicating stable docking of the barrel in close proximity to the tunnel exit. In contrast, NAC bearing mutations in the NACβ linker (HR:AA) or βh2 (KF:AA) more frequently sampled the medium and low FRET states (Fig. 3C). These differences are reflected in the FRET histograms, which show that WT NAC resides primarily in the high FRET state, whereas the FRET distribution for NAC switch mutants shifted to medium- and low-FRET states (Fig. 3D-E). These results demonstrate that the docking of the NAC barrel at the ribosome tunnel exit was impaired by mutations in the switch region.

We further measured the kinetic stability of NAC interactions with RNC_MTS_ using smTIRFM. Over 80% of WT NAC dissociated from RNC_MTS_ with a rate constant of ∼0.14 s^-1^, while the remainder dissociated at a faster rate (Extended Data Fig. 10A). The more-stably bound population (slow-dissociating) decreased to ∼40% and ∼20%, respectively, with NAC mutants HR:AA and KF:AA (Extended Data Fig. 10B). Thus, mutations in the NAC switch region impaired proper barrel docking on the ribosome surface, and the faster dissociation of these mutants compared to WT NAC are consistent with co-sedimentation assays showing reduced mutant binding to ribosomes (Fig. 2C-D). We note that despite the weakened binding, the association of HR:AA and KF:AA were still confidently detected under smTIRFM with < 5 nM NAC present, indicating that both NAC mutants still bind ribosomes with high affinity. Taken together, these data suggest that the molecular switch controls the conformational dynamics of the NAC barrel on the ribosome.

### MTS mutations destabilize the NAC barrel on the ribosome

We next asked whether the stabilized conformation of NAC barrel domain at the ribosome exit site is dependent on the presence of an MTS. To this end, we either deleted the amphiphilic helical segment of the MTS in the Oxa1L nascent chain (Oxa1L_ΛMTS_) or replaced this region with the ER signal sequence from a bona fide SRP substrate preprolactin (Oxa1L_MTS-to-SS_). smTIRFM experiments were performed to monitor the NAC barrel conformation on RNCs carrying the mutated MTS. Both mutations led to increased dynamics of the NAC barrel, substantially increasing the population in low and medium FRET states compared to that on RNC bearing the WT Oxa1L MTS (Figure 4A-C). In addition, the more stably bound (slow-dissociating) NAC population on these RNCs decreased to ∼40% and ∼60% for the Oxa1L_ΔMTS_ and Oxa1L_MTS-to-SS_ nascent chain, respectively, compared to 80% on Oxa1L_WT_ (Figure 4D-E).

**Figure 4.**
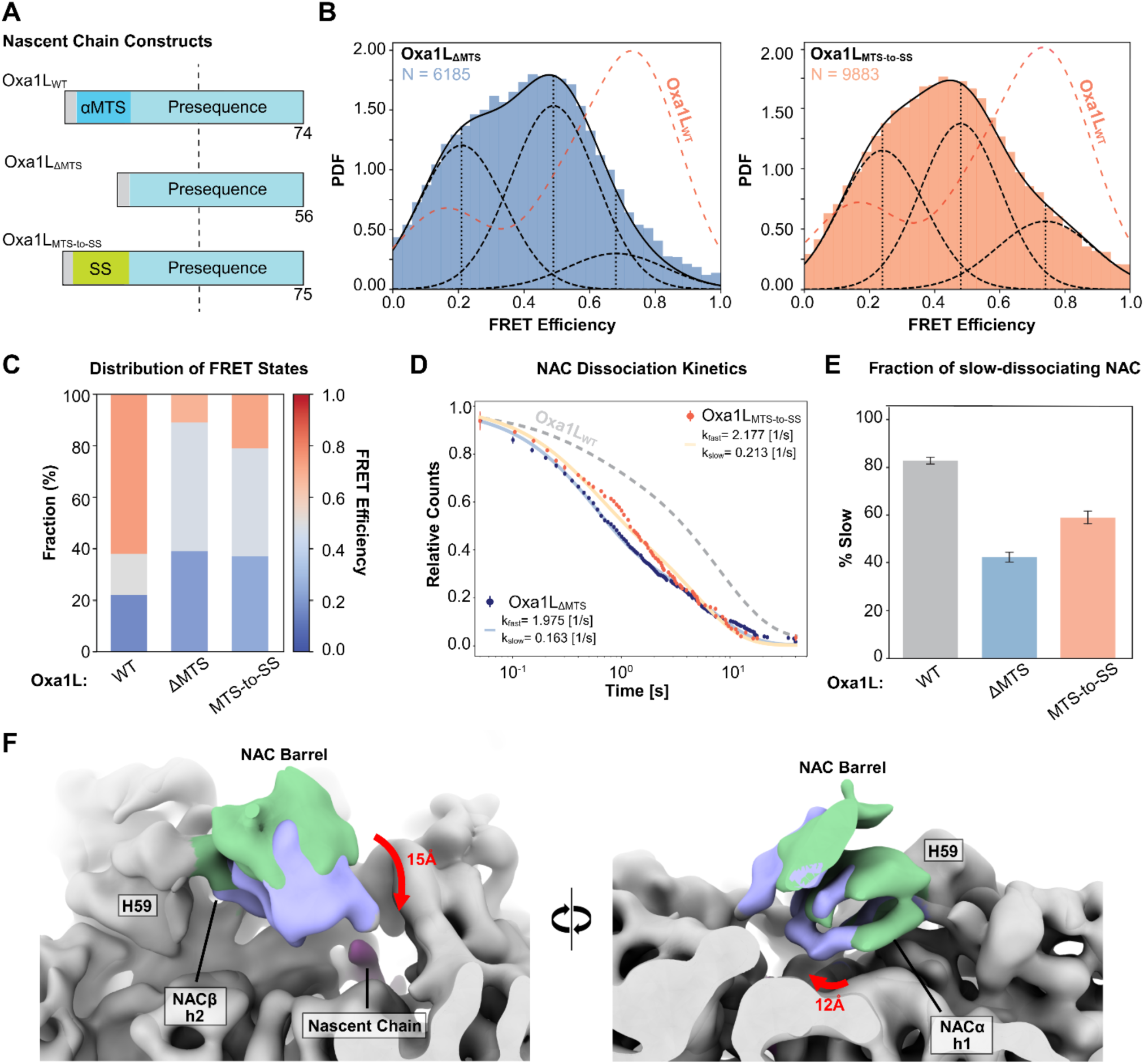
The MTS drives stable binding of the NAC barrel adjacent to the ribosome exit tunnel. **A**) Scheme showing the MTS mutations in the Oxa1L nascent chain. ‘αMTS’ (*dark blue*) denotes the amphiphilic N-terminal helix in the nascent chain, the rest of the presequence is in *light blue*. ‘SS’ denotes the hydrophobic core of the ER signal sequence from pPL. **B**) smFRET histograms for NAC bound to ribosomes displaying Oxa1L_ΔMTS_ (left panel) and Oxa1L_MTS-to-SS_ (right panel), using the same FRET pair as in Figure 3. The dashed line shows the data for wildtype Oxa1L RNCs for comparison. N, number of FRET events used to construct the histogram. PDF, probability density function. The data were fit to the sum (solid lines) of three Gaussian functions (dashed lines). **C**) Summary of the distribution of NAC in different FRET states, from analysis of the data in (B). **D**) Kinetics of NAC dissociation from the indicated RNCs. The lines show fits of the data to a double exponential function, which gave the indicated dissociation rate constants for the fast- (*k_fast_*) and slow- (*k_slow_*) dissociating populations. Grey dashed lines show the data with WT RNC_MTS_ for comparison. **E**) Summary of the fraction of slow-dissociating population of NAC bound to the indicated RNCs. From fits to the data in (D). Values are shown as fitted parameter ± fitting error. **F**) Close-up of the NAC barrel conformations on ribosomes translating Oxa1L_ΔMTS_. Cryo-EM maps filtered to 8Å are overlayed showing the NAC barrel in the upper (green) and shifted destabilized conformation (purple).

Based on the smFRET data, we hypothesized that mutation of the MTS destabilizes the NAC barrel, similar to previous structures of NAC bound to ribosomes displaying the nascent chain of an SRP substrate, preprolactin (pPL)^19^. We therefore determined the structure of NAC bound to ribosomes translating Oxa1L_ΔMTS_ (Extended Data Fig. 11A-B). Multiple NAC classes were resolved exhibiting barrel movements on the ribosome ranging from ∼12–15 Å (Figure 4F). Particles corresponding to the most extensive movement of the NAC barrel downward towards the PET (Extended Data Fig. 11C, ‘destabilized barrel state’) showed weak densities for the two amphipathic helices of NAC (Extended Data Fig. 12A). These destabilized conformations are reminiscent of previous structures of NAC on ribosomes translating an SRP substrate, which also displayed weaker densities for the amphipathic helices likely due to increased barrel dynamics (Extended Data Fig. 12B-C). Collectively, these results indicate that an MTS induces a stable conformation of the NAC barrel near the exit tunnel, which is likely crucial for preventing improper targeting of MTS-containing nascent chains by SRP.

### NAC mutants fail to regulate SRP conformation on RNCs

We next tested whether the new contacts made by the NAC switch region are important for preventing the mistargeting of ribosomes exposing a nascent MTS. SRP is activated for ER targeting when the SRP54 M-domain engages the ER signal sequence emerging from the exit tunnel, which induces SRP to adopt a ‘Proximal’ conformation in which the SRP54 NG-domain docks at uL23 and optimally associates with the SRP receptor at the ER^16,29,38^. Previously, we showed that NAC blocks SRP from adopting the targeting-active proximal conformation on ribosomes lacking an ER targeting signal and thus prevents mistargeting^16^. In the current conformation of NAC on RNC_MTS_, the NAC barrel is stabilized at the exit tunnel and would effectively block the binding of SRP54 M-domain at the exit tunnel (Fig. 5A), thus preventing the activation of SRP^16,27^. We hypothesize that the loss of the molecular switch would impair this sorting function of NAC, by reducing the stable docking of the NAC barrel at the polypeptide exit tunnel.

**Figure 5.**
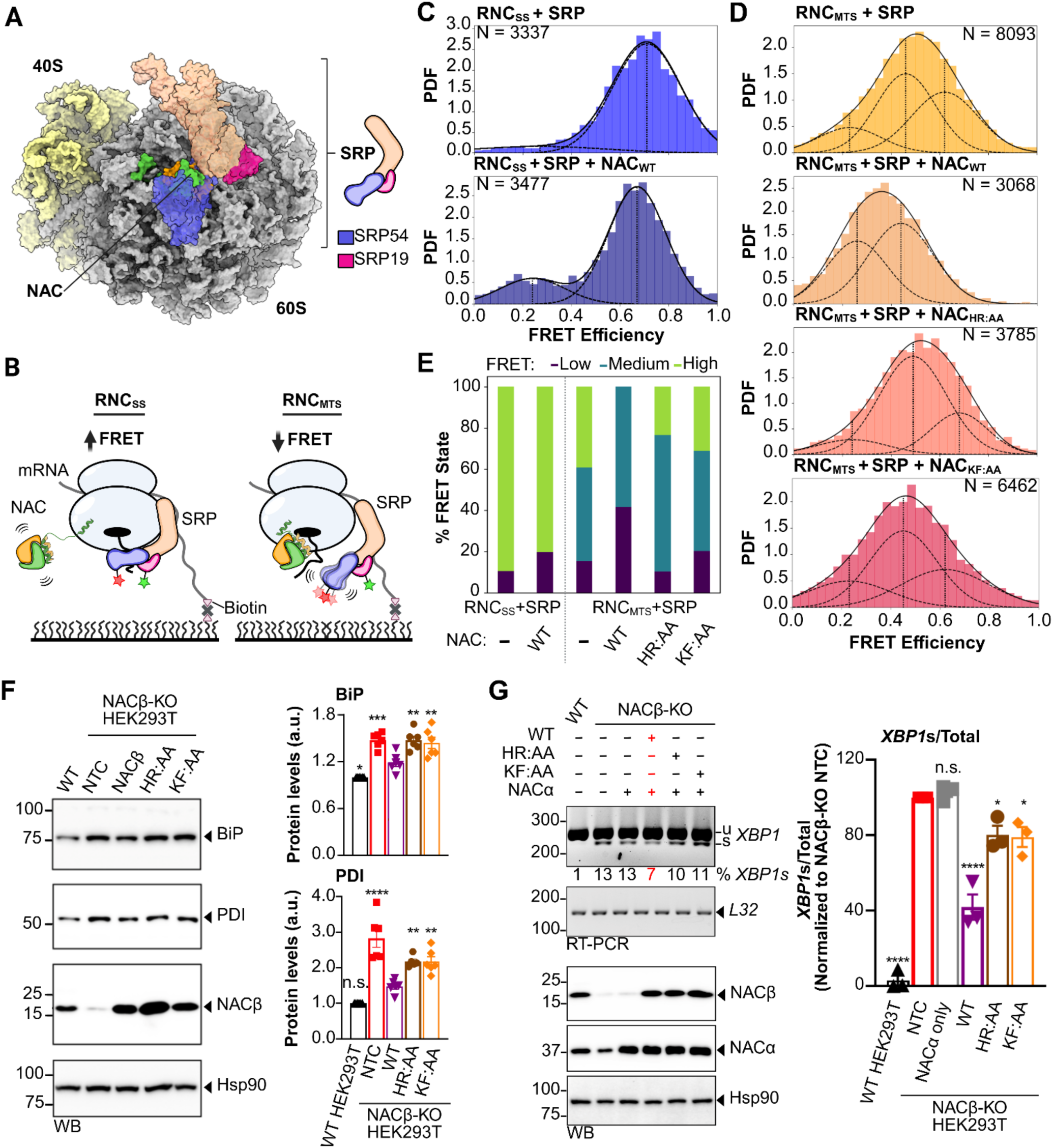
NAC molecular switch mutants fail to regulate SRP activation on the ribosome and lead to elevated ER-stress in human cells. **A)** Surface model depicting the overlap between SRP (PDB 7QWQ) and NAC (this study) on the ribosome. **B)** Scheme depicting the smTIRFM experiments to measure the conformational distribution of SRP on the ribosome. RNCs were immobilized on quartz slides. FRET was measured between a donor dye (green star) on SRP19 and an acceptor dye (red star) on the SRP54 NG domain. High FRET efficiency was observed when SRP is in the targeting active proximal conformation (left) and correlates with SRP-SRP receptor interaction rates^39^. **C, D)** smFRET histograms of SRP bound to ribosomes translating nascent chains containing an ER signal sequence (C) and an MTS (D), with or without the indicated NAC variant. PDF, probability density function. ‘N’ indicates the number of measurements. The data were fit to the sum (solid line) of two or three Gaussian functions (dashed lines), with the mean FRET value of each Gaussian indicated. **E)** Summary of the effect of WT and mutant NAC on the conformation of SRP bound to ribosomes translating ER (RNC_SS_) and mitochondrial nascent chains (RNC_MTS_). **F, G)** Experiments to measure ER stress in NAC knockout (KO) HEK293T cells, with or without transient overexpression of the indicated NAC variant. ER stress is measured by monitoring the expression of BiP and PDI protein (F) and levels of spliced (s) and unspliced (u) *XBP1* mRNA (G). Hsp90 and L32 are loading controls used for the WB and RT-PCR gels, respectively. The percentage of the ratio of spliced to total *XBP1* shown below the gel. The bar graphs in F and G show the quantification of the blots and gels from individual replicates normalized to controls. Values are plotted as mean ± SEM. Statistics: *p < 0.05, **p < 0.01, ***p < 0.001 and ****p < 0.0001 by one-way ANOVA followed by Dunnett’s multiple comparisons test. NTC-non-transfected control.

We conducted smTIRFM experiments to observe the conformational dynamics of SRP bound to ribosomes translating either an ER or a mitochondrial nascent chain. The proximal conformation of SRP was detected using FRET between a donor dye (Atto550) and an acceptor dye (Atto647N) labeled on SRP19 and SRP54, respectively (Fig. 5B)^39,40^. On ribosome bearing an ER signal sequence from preprolactin (pPL), SRP is dominated by a high FRET state that reports on the proximal conformation, both in the absence and presence of NAC (Fig. 5C), consistent with previous results^16^. On ribosomes exposing the MTS of Oxa1L, SRP is distributed between three conformations, with ∼40% of SRP in the high-FRET, proximal conformation that is targeting-active (Fig. 5D). The proximal conformation of SRP on RNC_MTS_ was abolished by WT NAC, whereas both NAC mutants failed to regulate the conformation of SRP (Fig. 5D-E). Thus, mutations in the switch region abolished the ability of NAC to prevent the mis-activation of SRP on ribosomes translating an MTS.

Previous work demonstrated that NAC knockdown led to the mistargeting of mitochondrial proteins to the ER, which induced modest ER stress and elevated expression of ER chaperones BiP and PDI^8^. We therefore tested the importance of the NAC molecular switch in preventing protein mistargeting to the ER by assessing the levels of ER chaperones and the splicing of the *XBP1* mRNA, both of which report on ER stress^41,42^. We first showed that NACβ KO HEK293T cells displayed increased levels of the ER chaperones, BiP and PDI (Extended Data Fig. 13A), as well as *XBP1* mRNA splicing (Extended Data Fig. 13B). We then reintroduced either WT or mutant NACβ in the NACβ KO cells. Overexpression of WT NACβ largely restored the ER stress markers to levels observed in WT cells, while both the NACβ HR:AA and KF:AA mutants failed to rescue the ER stress phenotype (Fig. 5F-G). Taken together, our results indicate that the NAC molecular switch prevents non-specific binding and activation of SRP to MTS-containing RNCs, and when perturbed, can lead to modest ER stress in human cells.

## Discussion

NAC is an essential protein biogenesis factor, conserved from yeast to humans, that interacts with all nascent chains^20,21^ as they emerge from the ribosome to facilitate proper protein sorting and biogenesis^3^. Based on our results, we propose that NAC adopts a stabilized conformation near the polypeptide exit tunnel when engaging ribosomes translating an MTS. This MTS-stabilized state of NAC is critical for preventing promiscuous SRP activation and ensures the correct sorting of mitochondrial proteins. In the absence of a complete MTS or the molecular switch that senses the MTS, the NAC barrel is more dynamic, allowing SRP to adopt a targeting-active conformation that can lead to mistargeting of nascent proteins (Figure 6).

**Figure 6.**
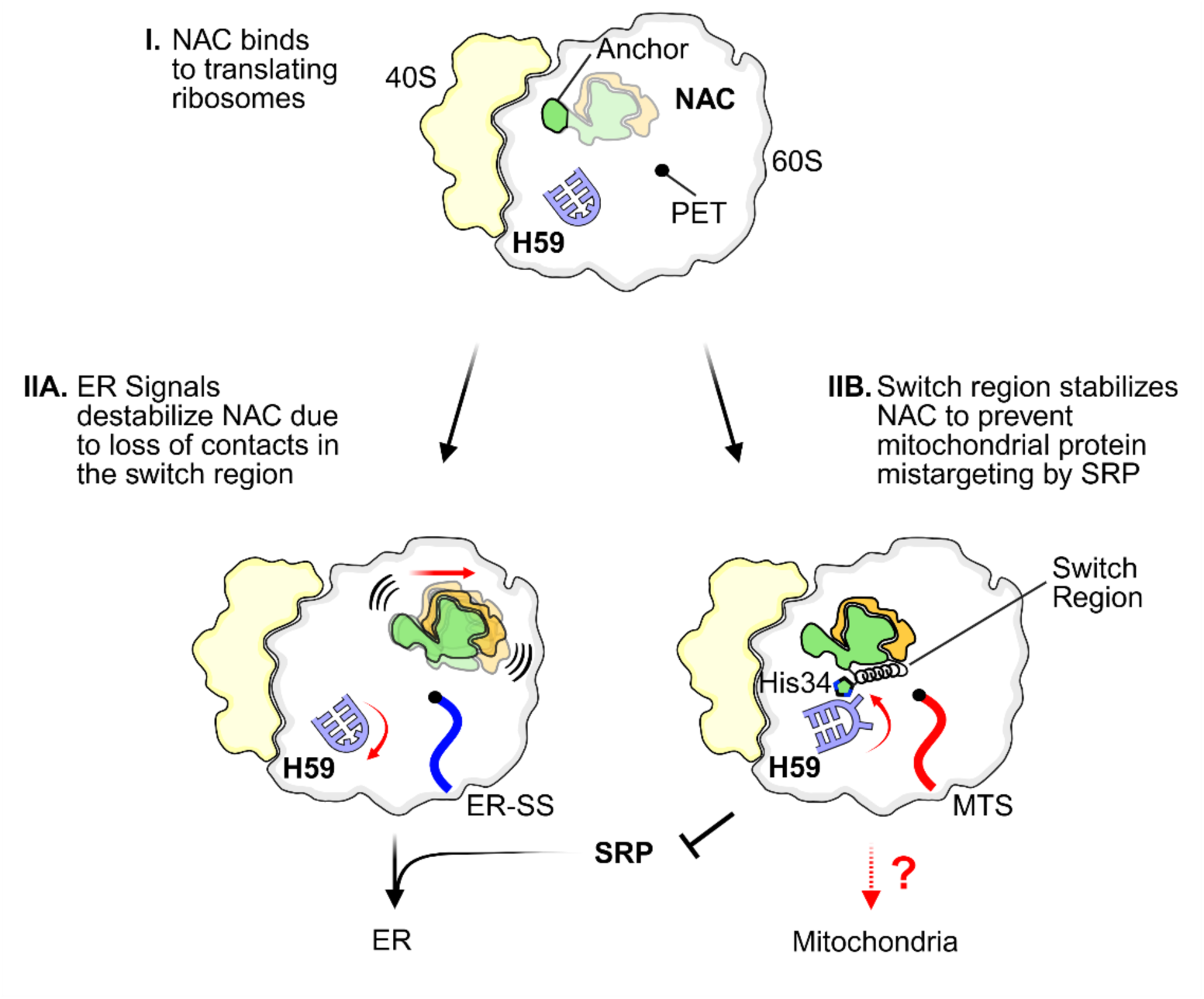
Model for MTS recognition by the NAC switch on translating ribosomes. NAC binds to all translating ribosomes using the NAC anchor (I). The NAC barrel domain can additionally dock near the polypeptide exit tunnel (PET), which is destabilized by hydrophobic ER signals, allowing SRP to engage the nascent chain and target the ribosome to the ER membrane (IIA). On MTS-containing ribosomes, the NAC molecular switch stabilizes the barrel domain near the PET accompanying a conformational change in H59. This prevents the mistargeting of mitochondrial proteins by SRP and enables their subsequent sorting to the mitochondria (IIB).

Recent studies showed that H59 interacts with NAC, SRP^19^, MAP1^23^, MAP2^43^, and the multi-functional protein EBP1^44^. Interestingly, the conformational change in H59 observed in the current structure and the subsequent repositioning of the barrel are different from a structure in which NAC coordinates the N-terminal processing of cytosolic nascent chains by methionine aminopeptidase 1 (MAP1) (Extended Data Fig. 14A)^23^. In the current conformation, both H59 and the NAC barrel clash with the position of MAP1 (Extended Data Fig. 14B-C). Thus, the conformational landscape of H59 may serve as an additional regulatory element to enhance the specificity of nascent chain sorting, processing, and targeting on the ribosome during translation.

Nascent chain properties such as hydrophobicity and secondary structural features are known to influence the interaction of NAC on the ribosome^16,19,20^. For example, hydrophobic ER signals weaken NAC binding on the ribosome, presumably by destabilizing the NAC barrel, to allow for SRP recruitment^19^. In contrast, both our structure and single-molecule experiments showed stable docking of the NAC barrel near the exit tunnel on ribosomes exposing an MTS. While we did not observe density for the MTS itself, likely due to the transient nature of NAC–nascent chain interactions^19,22,23^, our data provide strong evidence that NAC is able to sense an MTS early in translation. The presence of an MTS promotes the stabilized NAC conformation on the ribosome. Deletion of the characteristic amphiphilic helix in the MTS or substitution of this helical element with an ER signal sequence impairs the stable docking of the NAC barrel, reinforcing a direct role of the MTS in stabilizing NAC near the exit tunnel.

Mutagenesis experiments further demonstrate that the molecular switch plays a key role in mediating MTS sensing by stabilizing the new conformation of NAC on the ribosome, which is essential to prevent the non-specific recruitment and activation of SRP on ribosomes lacking an ER targeting signal^16^. Accordingly, NAC switch mutants fail to properly triage MTS-nascent chains, leading to the promiscuous activation of SRP on the ribosome. Furthermore, NAC mutants failed to fully rescue the ER stress caused by the depletion of NAC in human cells. Thus, stable docking at the ribosome exit by the NAC barrel domain is critical to prevent the mistargeting of mitochondrial nascent chains to the ER.

Our results provide new evidence for the role of NAC as a molecular gatekeeper at the ribosome. The toggling of the NAC barrel domain on ribosomes mediated by the newly discovered switch may allow NAC to sense newly synthesized proteins and contribute to mitochondrial protein sorting in humans. Disruption of this switch can lead to protein mistargeting and cellular stress, highlighting the importance of the regulatory mechanisms employed by NAC in maintaining cellular proteostasis. Furthermore, NAC enhances mitochondrial protein import and facilitates ribosome association with the mitochondrial surface^45–49^.

Our study reveals a critical molecular mechanism whereby NAC adopts a stabilized conformation on the ribosome in response to mitochondrial targeting sequences, shielding these nascent chains from promiscuous interactions with cytosolic factors and thereby promoting precise protein sorting and cellular proteostasis. Whether and how NAC facilitates ribosome recruitment to mitochondria remains an important unresolved question and requires future investigation.

## Materials and Methods

### Immunofluorescence

HEK293T cells were cultured on Millicell EZ SLIDE 8-well chamber (Millipore # PEZGS0816) for 24 hours. The cells were fixed by 4% paraformaldehyde (PFA) for 10 minutes at room temperature, followed by three washes with PBS. The fixed cells were simultaneously incubated with primary antibodies in blocking buffer (0.3% donkey serum and 0.25% Triton X-100 in PBS) overnight at 4 °C. Following three washes with PBS, the samples were incubated with secondary antibodies for 2 hours at room temperature. Mounting was performed with mounting medium containing DAPI (Vector Laboratories; #H-1200) and Fisherfinest Premium Cover Glasses (Fisher Scientific; 1#2-548-5P). Images were captured using the Zeiss LSM 980 confocal microscope at the University of Virginia Advanced Microscopy Facility. Antibodies for immunostaining were as follows: Anti-Hsp60 (mouse, Invitrogen #MA3-012, 1:200), anti-Tom20 (rabbit, Proteintech #11802-1, 1:200), anti-Oxa1L (rabbit, Proteintech #21055-1, 1:200), anti-Tom20 (mouse, Abcam #ab56783. 1:200). Secondary antibodies for fluorescent immunostaining (all 1:500) were as follows: Anti-rabbit IgG Alexa Fluor 488 (Jackson ImmunoResearch, #711-545-152), anti-Mouse IgG Alexa Fluor 555 (Invitrogen, #A32773).

### Protein Preparation

#### NAC Purification

A construct encoding for 6xHis-NACα and NACβ in a pET28b vector was expressed in *E. coli* BL21-CodonPlus DE3 competent cells (Agilent, cat no. 230245). QuickChange (Agilent, cat no. 210518) site-directed point mutagenesis was conducted to generate mutations in NACβ. The following procedures were used to purify all NAC variants. Cells were grown to OD 0.6-0.9 at 37 °C and induced with 1 mM IPTG at 18 °C overnight. Cells were pelleted, resuspended in 15 mL of lysis buffer (50 mM HEPES-KOH pH 7.5, 1M NaCl, 10% Glycerol, 6 mM β-mercaptoethanol, protease inhibitor (Roche, cOmplete mini, cat no.11836153001)), and lysed using a French press. The lysate was centrifuged using a TI 50.2 rotor at 20,000 rpm for 30 min at 4 °C (2X). The supernatant was loaded on a 5 mL HisTrap HP column (Cytiva, cat no. 17524802) using a P1 pump at 4 °C and washed with 1-2 column volumes (CV) of Nickel A buffer (50 mM HEPES-KOH pH 7.5, 1 M NaCl, 45 mM Imidazole, 10 % glycerol, 6 mM β-mercaptoethanol). The column was transferred to an Akta Pure fast protein liquid chromatography system (Cytiva) and washed for an additional 5 CV in nickel A buffer. Purified proteins bound to the column were eluted in a step gradient using 2-3CV of 15% and 30% nickel B buffer (50 mM HEPES-KOH pH 7.5, 150 mM KOAc, 300 mM Imidazole, 10 % glycerol, 6 mM β-mercaptoethanol).

Fractions from the 30% peak were pooled and added to 7K MWCO dialysis tubing (ThermoFisher, cat no. 68700) with 3C protease (Genescript, cat no. Z03092) to cleave the N-terminal His tag on NACα. Pooled fractions were dialyzed overnight at 4 °C in dialysis buffer (50 mM HEPES-KOH pH 7.5, 150 mM KOAc, 10% glycerol, 6 mM β-mercaptoethanol). Dialyzed proteins were loaded onto a HiTrap Q HP column (Cytiva, cat no. 17115401) using a P1 pump and the column was washed with 1-2 CV of ion exchange buffer A (50 mM HEPES-KOH, 2 mM DTT, 2 mM EDTA, 10% glycerol). The column was moved to the Akta and bound proteins were eluted from the column using a linear gradient of ion exchange buffer B to 50% (50 mM HEPES-KOH, 1 M NaCl, 2 mM DTT, 2 mM EDTA, 10% glycerol). Fractions eluted with∼30% ion exchange buffer B were combined and concentrated using an Amicon Ultra 4 mL centrifugal filter (Millipore, cat no. UFC801024). One volume of ion exchange buffer A was added to the sample and the concentration of NAC was determined on the Nanodrop (ThermoFisher) using a molar extinction coefficient, 2980 M^-1^ cm^-1^. The samples were snap frozen in liquid nitrogen and stored at −80 °C.

#### NAC labeling

WT NACα/NACβ(S57C) and ribosome-binding mutants were labeled with maleimide-Cy3B (Cytiva). The protein was exchanged into Labeling buffer (50 mM KHEPES, pH 7.5, 100 mM NaCl, 1 mM TCEP, 20% glycerol) and incubated with a 5-fold molar excess of dye for 2 hours at room temperature. Free dye was removed using a G-25 Sephadex size exclusion column (GE Healthcare). Fractions containing labeled NAC were identified by SDS-PAGE, pooled, and concentrated to 100-250 μM.

#### Purification and labeling of SRP subunits

Purification of SRP9/14, SRP19, SRP54, and SRP68/72 were described previously^39^. The single cysteine mutants SRP19(K64C) and SRP54(S12C) were purified as for WT proteins and labeled with maleimide-Atto550 and maleimide-Atto647N (ATTO-TEC), respectively, following published protocols^16,39^. Briefly, SRP19(K64C) and SRP54(S12C) were incubated with 2 mM DTT for 30 minutes at 25 °C. Reduced proteins were exchanged into the Labeling buffer by passing twice through a micro Bio-Spin column packed with Bio-Gel P-6 resin (Bio-Rad). Proteins were incubated with an 8-fold molar excess of maleimide dye for 3 hours at 4 °C. The labeling reaction was quenched with 2 mM DTT, and excess dye was removed using the G-25 Sephadex Fine size exclusion column (GE Healthcare) run in the labeling buffer (50 mM KHEPES, pH 7.5, 300 mM NaCl, 2 mM EDTA, 1 mM TCEP, 10% Glycerol). Fractions containing labeled SRP subunits were identified by SDS-PAGE, pooled, and concentrated to ∼40 μM.

#### SRP assembly

SRP was assembled as described^39^. Briefly, purified folded 7SL SRP RNA was incubated in the binding buffer (20 mM Tris, pH 7.5, 300 mM KOAc, 5 mM Mg(OAc)_2_, 5 mM DTT, 10% Glycerol) at 37 °C. Subsequently, the assembly reaction was performed in the HKMN buffer (50 mM KHEPES, pH 7.5, 500 mM KOAc, 5 mM Mg(OAc)_2_, 1 mM DTT, 0.02% Nikkol) with the sequential addition of the protein subunits in the following order: SRP19(K64C)-Atto550 for 10 minutes, SRP68/72 together with SRP9/14 for 10 min, and SRP54(S12C)-Atto647N for 30 minutes at 37 °C. Holo-SRP was purified on a DEAE-Sephadex anion exchange column (Sigma). Fractions containing fully assembled SRP were eluted in a buffer containing 600 mM KOAc, identified by A_260_ measurements, pooled, and stored at −80 °C. The activity of purified SRP was tested using a translocation assay with preprolactin as the SRP substrate, as described^16^.

### RNC Purification

#### PCR and in vitro Transcription

A construct encoding for Hsp60, Oxa1L_MTS_, or Oxa1L_ΔMTS_ was PCR amplified to include the upstream T7 promoter, 3X-FLAG tag, and the N-terminus of each nascent chain. PCR products were cleaned up using the QIAquick PCR Purification Kit (Qiagen, cat no. 28104). The resulting purified PCR products were *in vitro* transcribed for 4 hours at 38 °C, using 1.1 mg/mL T7 polymerase in reaction buffer (40 mM Tris-HCl pH 7.6, 5 mM ribonucleoside triphosphates, 6 mM MgCl_2_, 2 mM spermidine, 1 mM DTT, 0.04 U/uL RNase Inhibitor) or with the T7 HiScribe kit (NEB, cat no. E2040S). Following the reaction period, the precipitate was pelleted at 14,000 rpm for 5 minutes at 4 °C. The supernatant was transferred to a sterile 1.5 mL tube and 6M LiCl was added 1:1 for 1 hour on ice. After the incubation period, the mixture was centrifuged at 14,000 rpm for 20 minutes at 4 °C and the supernatant was discarded. The pellet was washed with 200 µL of ice cold 70% ethanol and centrifuged at 14,000 rpm for 5 minutes at 4°C. The supernatant was discarded and 100 µL of sterile ultrapure water was used to resuspend the pellet. The sample was placed on ice and 40 µL of 2.8 M NaOAc and 300 µL 95% ice cold ethanol were added. The mixture was incubated on ice for 5 minutes and centrifuged at 14,000 rpm for 30 minutes at 4 °C. The resulting pellet was washed with 200 µL of 70% ethanol and centrifuged at 14,000 rpm for 5 min at 4°C. The supernatant was discarded, and the resulting purified RNA was resuspended in 100 µL of sterile ultrapure water.

#### RNC purification following in vitro translation

Rabbit reticulocyte lysates (Promega, cat no. L4540) were reacted with Hsp60 or Oxa1L RNA after diluting the lysate to 66.7% with buffer (RNase Inhibitor (Promega) 0.04 U/µL, 0.5X protease inhibitor (Promega), 24 µM amino acid mix, 81 mM KCl, 2 mM Mg(OAc)_2_). The reaction was incubated at 32 °C for 25 minutes and then placed on ice.

A 1 mL polypropylene gravity column (Qiagen, cat no. 34924) was loaded with Anti-DYKDDDK G1 affinity resin (GenScript, cat no. L00432 or Sigma, cat no. A2220). The column was rinsed with 10 bead volumes of 1X PBS followed by 10 bead volumes of low-salt wash buffer (50 mM HEPES-KOH pH 7.7, 100 mM KCl, 10 mM MgCl_2_). The RRL reaction was added to the column and incubated for 2 hours at 4 °C with rotation. After the incubation period, the cap and side walls of the gravity column were rinsed with high-salt wash buffer (50 mM HEPES-KOH pH 7.7, 750 mM KOAc, 10 mM Mg(OAc)_2_, 0.1% Triton-X, 1 mM DTT). The reaction supernatant was run through the column and the beads were subsequently washed with 10 bead volumes of high-salt wash buffer (2X) followed by 10 bead volumes of low-salt wash buffer (2X). RNCs were incubated with elution buffer (50 mM HEPES-KOH pH 7.7, 100 mM KOAc, 15 mM Mg(OAc)_2_) containing 0.25 mg/mL FLAG peptide for 15 minutes at RT (5X). The eluted fractions were placed on ice and combined to a total volume of 500 µL and centrifuged at 100, 000 rpm for 1 hr at 4 °C using the TLA120.1 rotor (Beckman). The pellet was resuspended in base elution buffer by gently pipetting up and down. The ribosome concentrations were determined using RNA reading at A260 on the Nanodrop. RNCs were snap frozen in liquid nitrogen and stored at −80 °C.

#### RNC labelling and purification for smTIRFM

*In vitro* translations used a pUC19 vector containing, from 5’ to 3’, a T7 promoter, encephalomyocarditis virus (EMCV) internal ribosome entry site (IRES), 3xFLAG-tag, and 3C protease site19. DNA fragment encoding the nascent chain of preprolactin (pPL) or Oxa1L was cloned after the 3C protease site using Gibson assembly. In the Oxa1L_ΔMTS_ construct, an amphipathic helical region (MTS) of the Oxa1L presequence (LMCGRRELLRLLQSGRRV, residues 4-21) was deleted, whereas in the Oxa1L_MTS-to-SS_ the deleted region was swapped with signal sequence of preprolactin (LLLLLLVSNLLL, residues 13-24). For fluorescent labeling of the RNCs, an amber codon was introduced at residue 39 in the Oxa1L nascent chain using QuickChange mutagenesis.

PCR fragments encoding the region from the T7 promoter to Ser74 of Oxa1L or Phe80 of pPL were transcribed using the T7 MegaScript protocol in the presence of 5 mM 5’-biotin-G-monophosphate (TriLink). The resulting mRNAs were translated in rabbit reticulocyte lysate (RRL, Green Hectares) supplemented with 1 µM *Methanosarcina mazei* pyrrolysine synthase (*Mm*PylRS), 10 mg/L *M. mazei* amber suppressor tRNA (*Mm*PyltRNA), and 100 µM axial-trans-cyclooct-2-en-L-450 lysine (TCOK, SiChem), for 30 minutes at 32 °C. The preparation of *Mm*PylRS and *Mm*PyltRNA was described in Hsieh *et al*^16^.

The translation reaction was layered on a High Salt Sucrose Cushion (50 mM KHEPES, pH 7.5, 1 M KOAc, 15 mM Mg(OAc)_2_, 0.5 M Sucrose, 0.1% Triton, 2 mM DTT) at a volumetric ratio of 2:3. The ribosomal fraction was pelleted by ultracentrifugation (100k rpm for 45 minutes at 4°C) in a TLA100.3 rotor (Beckman Coulter). Pellets were resuspended in RNC buffer (50 mM KHEPES, pH 7.5, 150 mM KOAc, 2 mM Mg(OAc)_2_) to 1 µM ribosome concentration and incubated with 1 µM tetrazine-conjugated Atto647N dye (Jena Biosciences) for 30 minutes at 25 °C, to allow for the Diels-Alder cycloaddition reaction. Labeled RNCs were incubated with anti-DYKDDDK magnetic agarose (Pierce) pre-equilibrated in RNC buffer for 1 hour at 4 °C with constant rotation. Beads were washed sequentially with 10 bead volumes of RNC buffer containing 300 mM KOAc, RNC buffer containing 0.1% Triton, and RNC buffer. RNC was eluted with 1.5 mg/ml 3xFLAG (DYKDDDK) peptide for 30 minutes at 4 °C with constant rotation. The eluted RNCs were incubated with 1 µM 3C protease (GoldBio) for at least 2 hours at 25°C, and sedimented using a High Salt Sucrose Cushion at a volumetric ratio of 2:3 at 100k rpm for 30 minutes at 4 °C in a TLA120.2 rotor (Beckman Coulter). Pelleted RNC was resuspended in storage buffer (50 mM KHEPES, pH 7.5, 150 mM KOAc, 5 mM Mg(OAc)_2_, 0.04% NIKKOL, 2 mM DTT) to ∼100 nM (ribosome concentration). The presence of fluorescently labeled nascent chain was confirmed by SDS-PAGE and fluorescence scanning for Atto674N signal (Typhoon biomolecular imager, Cytiva). Fluorescently labeled RNCs were snap frozen in liquid nitrogen and stored at −80 °C.

#### Single-molecule TIRF microscopy

Surfaces of pure quartz imaging slides and coverslips were amminosilanized with vectabond (Vector lab) and coated with biotin-PEG (Laysan Bio). Slides were passivated with passivation buffer (1X Tween-20, 1 mg/ml BSA) for at least 1 hour at 25 °C, washed with 50 mM KHEPES, pH 7.5, and incubated with 0.5 mg/ml NeutrAvidin (ThermoFisher) for 10 minutes. Unbound NeutrAvidin was washed away with Imaging Buffer, which constitutes Assay Buffer (50 mM KHEPES, pH 7.5, 150 mM KOAc, 5 mM Mg(OAc)_2_, 0.04% NIKKOL, 2 mM DTT) supplemented with 1 mg/ml BSA, 4 mM Trolox, 2.5 mM protocatechuic acid (PCA), and 50 nM protocatechuate-3,4-dioxygenase from *Pseudomonas sp.* (PCD, Sigma)^50^. For assays involving SRP, the imaging buffer further included 200 μM non-hydrolyzable GTP analog, guanosine-5′-[(β,γ)-imido]triphosphate (GppNHp).

RNC with 3’-biotinylated mRNA was diluted to 1.5 nM in Imaging Buffer and immobilized on NeutrAvidin-coupled slides for 10 minutes at 25 °C. For experiments with NAC, RNC-coated slides were then flushed with 2 nM WT or mutant Cy3B-labeled NAC in Imaging Buffer. For experiments involving NAC and SRP, 1.5 nM doubly labeled SRP and 2 nM unlabeled WT or mutant NAC were incubated in Imaging Buffer for 5 minutes and loaded onto slides with immobilized RNCs.

Movies were recorded using MicroManager on a custom-built TIRF microscopy setup^51^. The presence of immobilized Atto647N-conjugated RNCs was confirmed by excitation at 635 nm. Movies were recorded by excitation at 532 nm (donor dye; Atto550 or Cy3B) in single-excitation mode and detection of both the donor and acceptor channels, with a temporal resolution of 50 ms.

#### Analysis of smTIRF microscopy data

Donor and acceptor channel image series were aligned and analyzed using iSMS software^52^. FRET efficiencies were calculated based on the raw intensity of donor and acceptor fluorescence time traces and were corrected for background noise and 𝛾 factor, which accounts for the difference in quantum yield between the dyes and leakage of donor fluorescence to the acceptor channel. Apparent FRET efficiency (*E*_app_) was calculated using Equation 1,

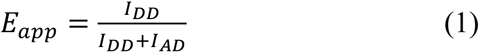

in which 𝐼_𝐷𝐷_ and 𝐼_𝐴𝐷_ are the fluorescence intensities of the donor and acceptor dye, respectively, upon excitation of the donor. FRET traces from all movies of the same sample were combined and analyzed using Hidden Markov Modelling (HMM) available in the iSMS software. The number of FRET states was established by applying Bayesian information criterion. These analyses yielded three FRET populations and the mean FRET efficiency for each state. These parameters were used for Gaussian fitting of the FRET histograms according to Eq 2,

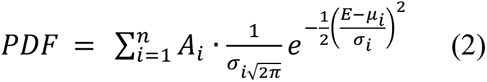

in which PDF is the probability density function, 𝑛 is the number of FRET states detected and is set to 3 (determined by HMM), 𝐴_𝑖_ is the weight of the ith Gaussian, and 𝜎_𝑖_ and 𝜇_𝑖_ are the standard deviation and center (determined by HMM) of the ith Gaussian.

To determine the residence time of NAC on RNC, cumulative fluorescence intensity signal (*I*_cumulative_) was calculated using Eq 3,

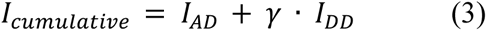

and plotted over time for each trace. Colocalization of labeled NAC and RNC identified by 𝐼_𝑐𝑢𝑚𝑢𝑙𝑎𝑡𝑖𝑣𝑒_ over background noise. Two-state HMM was then used to distinguish bound versus dissociated NAC and determine the dwell time of each colocalization event (*t*). The cumulative probability distribution of *t* was fitted to Eq 4,

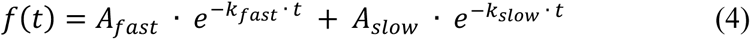

in which *A_fast_* and *A_slow_* are the amplitudes of the fast and slow-dissociating populations, and *k_fast_* and *k_slow_*are the respective dissociation rate constants.

### Cryo-EM sample preparation and data acquisition

FLAG-SUMO-Hsp60 RNCs (∼350 ng/uL) were reacted with 260 nM SUMO protease (Trialtus, cat no 30-1030) at 30°C for 20 minutes. NAC was added at a final concentration of 2 µM in reaction buffer (50 mM HEPES-KOH pH 7.7, 100 mM KOAc, 15 mM Mg(OAc)_2_, 0.01875% GDN) for 20 minutes at 30°C. The reaction was moved to ice prior to freezing grids. Quantifoil R2/1 300 mesh copper grids (Quantifoil, cat no. Q350CR1) were freshly coated with 3.3 nm carbon and glow discharged at 15 mA for 15 seconds using an EMS glow discharger. A 5 µL drop of the NAC-RNC reaction was incubated for 1 minute on the grid at 100% relative humidity and 4°C before blotting. The grid was subsequently plunge frozen into a cup filled with liquid ethane pre-cooled to liquid nitrogen temperatures. Cryo-EM data was collected on the ThermoFisher Glacios electron microscope operated at 200 kV and equipped with a Falcon4 direct electron detector at a total magnification of 120, 000x using a defocus range of −2.2 to −1.2 and a step size of 0.2. Movies were collected at a pixel size of 1.2 Å with a dose of 50 e/Å^2^ across 40 frames per movie.

FLAG-3C-Oxa1L_MTS_ and FLAG-3C-Oxa1L_ΔMTS_ RNCs (∼200-250 ng/uL) were reacted with 250 nM 3C protease at 30°C for 20 minutes. NAC was added at a final concentration of 1 µM in reaction buffer (50 mM HEPES-KOH pH 7.7, 100 mM KOAc, 15 mM Mg(OAc)_2_, 0.02% C12E8) for 20 minutes at 30°C. The reaction was moved to ice prior to freezing grids. Quantifoil R2/1 300 mesh copper grids (Quantifoil, cat no. Q350CR1) were freshly coated with 3.3 nm carbon and glow discharged at 15 mA for 15 seconds using an EMS glow discharger. A 5 µL drop of the NAC-RNC reaction was incubated for 1 minute on the grid at 100% relative humidity and 4°C before blotting. The grid was subsequently plunge frozen into a cup filled with liquid ethane pre-cooled to liquid nitrogen temperatures. Cryo-EM data was collected on the ThermoFisher Krios electron microscope operated at 300 kV and equipped with a K3 direct electron detector at a total magnification of 105, 000x with an energy filter slit width of 10 eV, using a defocus range of −1.8 to −0.8 with a step size of 0.2. Movies were collected at a pixel size of 0.83 Å with a dose of 32 e/Å^2^ (Oxa1L_MTS_) or 40 e/Å^2^ (Oxa1L_ΔMTS_) across 40 frames per movie.

### Cryo-EM data processing strategy

#### Hsp60 RNC_MTS_

Movies were motion-corrected, dose-weighted, and the contrast transfer function (CTF) was estimated in CryoSPARC^53,54^. Template picker was used to pick particles with a diameter of 250 Å from the resulting micrographs. A total of 1,366,067 particles were extracted from 8,780 micrographs at a box size of 512 x 512 pixels. The extracted particles underwent 2D classification, with ribosome classes selected for further processing. A homogenous refinement of the selected 519,343 particles yielded a consensus 80S ribosome map at 2.97 Å. Following refinement, translating ribosomes with P-site tRNA density were classified using a spherical mask on the A/P/E by conducting a 3D variability analysis in CryoSPARC^55^. Particles with a stable P-site tRNA density were refined separately and subjected to an additional 3D variability analysis, using a spherical mask on the exit tunnel to identify particles with strong NAC density. An additional 3D variability analysis was conducted on the exit tunnel to remove particles with noisy artifacts. The final set of 36, 986 particles with stable NAC and P-site tRNA density were polished using the reference-based motion correction package implemented in CryoSPARC. A homogenous refinement was conducted on the polished particles which yielded the Hsp60 NAC-RNC_MTS_ map at a global resolution of 3.04 Å-determined by the gold standard Fourier Shell Correlation (FSC) of two half-sets of particles processed independently with an FSC threshold of 0.143. The local resolution of the map was calculated in CryoSPARC at an FSC threshold of 0.143.

#### Oxa1L RNC_MTS_ and Oxa1L RNC_ΔMTS_

Both Oxa1L RNC MTS and Oxa1L RNC ΔMTS were processed in the same manner. Movies were motion and CTF corrected in CryoSPARC. Blob picker was used to pick particles with a diameter of 250-400 Å. Picked particles were extracted at a box size of 512 x 512 pixels and binned to 128 x 128 pixels for 2D classification. Ab-initio reconstruction was used to generate reference maps for 3D classification of all extracted particles to select 80S ribosome particles and exclude poorly aligning or junk particles. A homogenous refinement was conducted on the final set of 80S ribosome particles. Subsequent 3D variability analyses were conducted to sort for particles with stable P-site tRNA and NAC density using focused spherical masks. Particles were reextracted at full pixel size and refined yielding high-resolution maps of ribosomes translating the Oxa1L MTS and the MTS deletion mutant of Oxa1L. The local resolution was calculated in CryoSPARC at an FSC threshold of 0.143.

### Model Building

Following data processing, a model of the 80S ribosome (PDB 6R5Q) and NAC (PDB 7QWR) was docked into the cryo-EM map using ChimeraX. NAC and eL22 were further fitted into the map as rigid bodies in Coot and manually adjusted based on side chain densities observed in the map. The unresolved N-terminal fluke of NACβ was built *de novo* guided by the visible side chain density. PHENIX was used to refine the model in the NAC-RNC_MTS_ density map with 3 macrocycles of real space refinements applying Ramachandran, sidechain rotamer, and protein secondary structure to correct for clashes. The final model was validated using MolProbity in PHENIX and figures were made in ChimeraX.

### Biochemistry Experiments

#### Isolation of Ribosomes from HEK293 Cells

Ribosomes were isolated from HEK293 GnTI cell pellets. Cell pellets were resuspended in lysis buffer (50 mM HEPES-KOH pH 7.7, 100 mM KOAc, 5 mM Mg(OAc)_2_, 0.5% IGEPAL, 1 mM DTT, 0.5X Protease inhibitor (Promega), 0.04 U/uL RNase inhibitor (Promega)) and lysed by passing the resuspension through a 23G x 1 ¼ needle four times (BD, cat no. 305120). The lysate was centrifuged at 11, 000 xg for 10 min at 4 °C. The supernatant was layered 1:1 on a sucrose cushion (50 mM HEPES-KOH pH 7.7, 750 KOAc, 5 mM Mg(OAc)_2_, 2 mM TCEP, 30% sucrose (w/v)) and centrifuged at 100, 000 rpm for 1 hr at 4°C using a TLA 120.1 rotor. The pellets were resuspended in ribosome buffer (50 mM HEPES-KOH pH 7.7, 100 mM KOAc, 2 Mg(OAc)_2_, 2 mM TCEP, 0.5X protease inhibitor, 0.04 U/uL RNase inhibitor (Promega)). The concentration was determined on the Nanodrop (ThermoFisher) and aliquots of isolated ribosomes were snap frozen in liquid nitrogen.

#### NAC-Ribosome Co-sedimentation Assay

NAC variants and ribosomes were thawed on ice. Ribosomes were centrifuged at 13,000 rpm for 10 min at 4°C and the supernatant was transferred to a sterile 1.5 mL tube. NAC was diluted to 10 µM in reaction buffer (50 mM HEPES-KOH pH 7.7, 100 mM KOAc, 15 mM Mg(OAc)_2_, 0.04 U/uL RNase Inhibitor (Promega)). Ribosomes were diluted to 13 A260 units/mL in reaction buffer and NAC variants were added at a final concentration of 0.2 µM. The reaction was incubated at 30°C for 20 minutes and then moved to ice. A 50 µL portion of the reaction was layered on top of a sucrose cushion (50 mM HEPES-KOH pH 7.7, 100 mM KOAc, 15 mM Mg(OAc)_2_, 0.5X protease inhibitor (Promega), 0.04 U/µL RNase inhibitor (Promega), 25% sucrose (w/v)) and centrifuged at 100, 000 rpm for 1 hr at 4°C. The pellet was resuspended in 1X SDS-PAGE loading dye (50 mM Tris-HCl pH 6.8, 2% SDS, 1% β-mercaptoethanol, 6% glycerol, 0.004% bromophenol blue) for analysis by Western Blot.

#### Western Blot

Samples analyzed by WB were prepared in 1X SDS-PAGE sample buffer (50 mM Tris-HCl pH 6.8, 2% SDS, 1% β-mercaptoethanol, 6% glycerol, 0.004% bromophenol blue). Proteins were resolved on a SurePAGE (Genscript, cat no. M00653, M00654) or NuPAGE (ThermoFisher, cat no. NP0323BOX) 4-12% Bis-Tris minigel using MES buffer (Genscript, cat no. M00677) and transferred onto a 0.2 µm nitrocellulose membrane (LI-COR, cat no. 926-31092). The membrane was blocked in 5% milk in 1X PBST (0.1% Tween-20) for 1hr at RT. After blocking, antibodies diluted in 2% BSA in 1X PBST were incubated with membranes. Primary antibodies used: FLAG (Sigma, cat no. F1804), NACβ (Invitrogen, cat no. PA5-63299), RPL10a (Invitrogen, cat no. MA5-44710). Secondary antibodies used: anti-mouse (Invitrogen, cat no. A21058) and anti-rabbit (Invitrogen, cat no. A32735). The LI-COR Odyssey imager was used for detection.

### NAC Rescue Experiments in HEK293T cells

#### CRISPR/Cas9-base Knockout HEK293T cells

HEK293T cells, obtained from ATCC, were cultured at 37°C with 5% CO2 in DMEM with 10% fetal bovine serum (Fisher Scientific). Generation of NACβ KO HEK293T cells was performed using CRISPR/Cas9. sgRNA oligonucleotides designed for human NACβ (5’-TGCTCGCAGAAAGAAGAAGG-3’) was inserted into lentiCRISPR, version 2 (Addgene 52961). Cells grown in 10 cm petri dishes were transfected with indicated plasmids using 5 μl 1 mg/ml polyethylenimine (PEI) (Millipore Sigma) per 1 μg of plasmids for HEK293T cells. The cells were cultured 24 hours after transfection in medium containing 2 μg/ml puromycin for 24 hours and then in normal growth medium. Single cell isolations were conducted to select for HEK293T cells with NACβ knocked out.

#### Western blot and antibodies

HEK293T cells were harvested and snap-frozen in liquid nitrogen. The proteins were extracted by sonication in NP-40 lysis buffer (50 mM Tris-HCl at pH7.5, 150 mM NaCl, 1% NP-40, 1 mM EDTA) with protease inhibitor (Millipore Sigma), DTT (Millipore Sigma, 1 mM), and phosphatase inhibitor cocktail (Millipore Sigma). Lysates were incubated on ice for 30 minutes and centrifuged at 16,000g for 10 minutes. Supernatants were collected and analyzed for protein concentration using Bio-Rad Protein Assay Dye (Bio-Rad). From 10 to 30 μg of protein was denatured at 95°C for 5 minutes in 5× SDS sample buffer (250 mM Tris-HCl pH 6.8, 10% sodium dodecyl sulfate, 0.05% bromophenol blue, 50% glycerol, and 1.44 M β-mercaptoethanol). Protein was separated using SDS-PAGE, followed by electrophoretic transfer to PVDF (Fisher Scientific) membrane. The blots were incubated in 2% BSA/TBST with the following primary antibodies overnight at 4°C: anti-HSP90 (Santa Cruz Biotechnology Inc., sc-13119, 1:5,000), anti-BiP (Abcam, #ab21685, 1:5000), anti-PDI (Enzo, #ADI-SPA-890, 1:5000), anti-NACα (Biorbyt, orb411671, 1:1000). Membranes were washed with TBST and incubated with HRP-conjugated secondary antibodies (Bio-Rad, 1:10,000) at room temperature for 1 hour for ECL Chemiluminescence Detection System (Bio-Rad) development. Band intensity was determined using Image Lab (Bio-Rad) software, version 6.1.

NACβ *Overexpression*

NACβ KO HEK293T cells grown in 3.5 cm petri dishes were transfected with 1 μg indicated plasmids with 5 μl 1mg/ml Polyethylenimine “Max” (PEI MAX) (Polysciences, 24765). The cells were cultured 24 hours after transfection and then harvested for Western blot.

#### RNA preparation and Reverse Transcription PCR (RT-PCR)

*XBP1* splicing was assessed in HEK293T cells as described previously^56,57^. Briefly, total RNA was extracted from cells using TRI Reagent and BCP phase separation reagent (Molecular Research Center, TR 118). RT-PCR primer sequences are:

*hXBP1*: F: 5’-GAATGAAGTGAGGCCAGTGG-3’ R: ACTGGGTCCTTCTGGGTAGA

*hL32* F: 5’-AGTTCCTGGTCCACAACGTC-3’ R: 5’-TTGGGGTTGGTGACTCTGAT

“h” denotes human genes.

#### Thapsigargin (TG) treatment

HEK293T cells treated with 100 nM thapsigargin for 4 hours were included as positive controls for ER stress.

## Acknowledgements

We thank the Jomaa, Shan, and Qi lab members for the helpful discussions. Cryo-EM data collection was conducted at the molecular electron microscopy core facility (RRID:SCR_019031) at the University of Virginia (UVA) School of Medicine which was built with NIH grant G20-RR31199. We thank Michael Purdy for assisting with the cryo-EM data collection and additional computational support. We thank Sarah Marks, Kinga Malezyna, and Travis Bishop for assisting with ribosome isolations, NAC purifications, and other biochemical assays. We also thank Shuangcheng Alivia Wu for initial help with preparation of NACβ KO cells. This work was supported by The Owens Family Foundation, by the Searle Scholars Program, Grant #: SSP-2023-106, and aided by Grant # 134088-IRG-19-143-33-IRG from the American Cancer Society to A.J., by National Institutes of Health grant R35 GM136321 and National Science Foundation grant 2219287 to S.S., by National Institutes of Health grant R01DK120047, R01DK120330, R35GM130292 to L.Q. We acknowledge the cellular and molecular biology training program at UVA for support provided to E.M. through NIH T32GM139787-3 and the medical scientist training program for support provided to Z.J.L. In addition, the National Ataxia Foundation is acknowledged for support provided to L.L.L and Z.J.L through Post- and Pre-doctoral Fellowships NAF 918037 and 1036307, respectively.

## Author Contributions

This study was conceived by A.J., S.S., E.M. and L.Q. Ribosomes were purified by E.M. and R.G. for structural and single molecule experiments, respectively. Y.P. provided materials used in various biochemical experiments throughout the study (site directed point mutagenesis, plasmid preps, cultured cells for ribosome isolations and assisted with the NAC purifications). E.M. collected, processed cryo-EM data, and built the atomic model and assembled the structural snapshots. R.G. conducted the single molecule total internal reflection microscopy studies. L.L.L., L.E.Z., Z.J.L., and S.A.W. generated the NACβ KO cells and conducted the cell biology experiments. A.J., S.S., and L.Q. supervised the work. E.M., A.J., S.S., and R.G. wrote the manuscript. All authors contributed to data analysis and the final version of the manuscript.

## Data Availability

The cryo-EM maps and corresponding atomic models have been deposited with the following accession codes: EMD-48552 and PDB-9MR4 for the Oxa1L NAC-RNC_MTS_ structure; EMD-71310 for Hsp60; EMD-71286 and EMD-71287 for the destabilized and lifted barrel states of the Oxa1L_ΔMTS_ structure, respectively. All data related to this study are available in the main text, extended data, or the supplementary material.

