## Extended Data Figures and Tables for "A molecular switch in NAC prevents mitochondrial protein mistargeting by SRP"

Maldosevic et al.

**Extended Data Contents:**

**Extended Data Figure 1.** Mitochondrial proteins are mislocalized in NACβ-KO HEK293T cells.
**Extended Data Figure 2.** Purification of the NAC complex and RNCs.
**Extended Data Figure 3.** Processing scheme for the structure of NAC in complex with ribosomes translating the N-terminal MTS of Oxa1L.
**Extended Data Figure 4.** Processing strategy and focused 3D classification scheme of NAC in complex with ribosomes translating the N-terminal MTS of Hsp60.
**Extended Data Figure 5.** Comparison of the Hsp60 and Oxa1L $RNC_{MTS}$ maps.
**Extended Data Figure 6.** Nascent chain density for the Oxa1L NAC-$RNC_{MTS}$ complex.
**Extended Data Figure 7.** Structure of the NACβ N-terminal fluke and anchor domain.
**Extended Data Figure 8.** Interactions of the NAC amphipathic helices with the ribosome.
**Extended Data Figure 9.** NACβ switch mutants influence NAC binding to the ribosome.
**Extended Data Figure 10.** NACβ switch mutants exhibit reduced kinetic stability on $RNC_{MTS}$.
**Extended Data Figure 11.** Processing strategy and focused 3D classification scheme of NAC in complex with ribosomes translating the MTS deletion mutant of Oxa1L.
**Extended Data Figure 12.** Map and model comparison of NAC-$RNC_{MTS}$ to previous NAC structures on ribosomes translating $SS_{pPL}$ and tubulin.
**Extended Data Figure 13.** NACβ knockout induces modest ER stress that elevates ER resident chaperones and increases the splicing of *XBP1* mRNA.
**Extended Data Figure 14.** Comparison of the NAC-$RNC_{MTS}$ structure with the previous MAP and EBP1 structures.
**Extended Data Table 1.** Cryo-EM data collection, refinement and validation statistics.

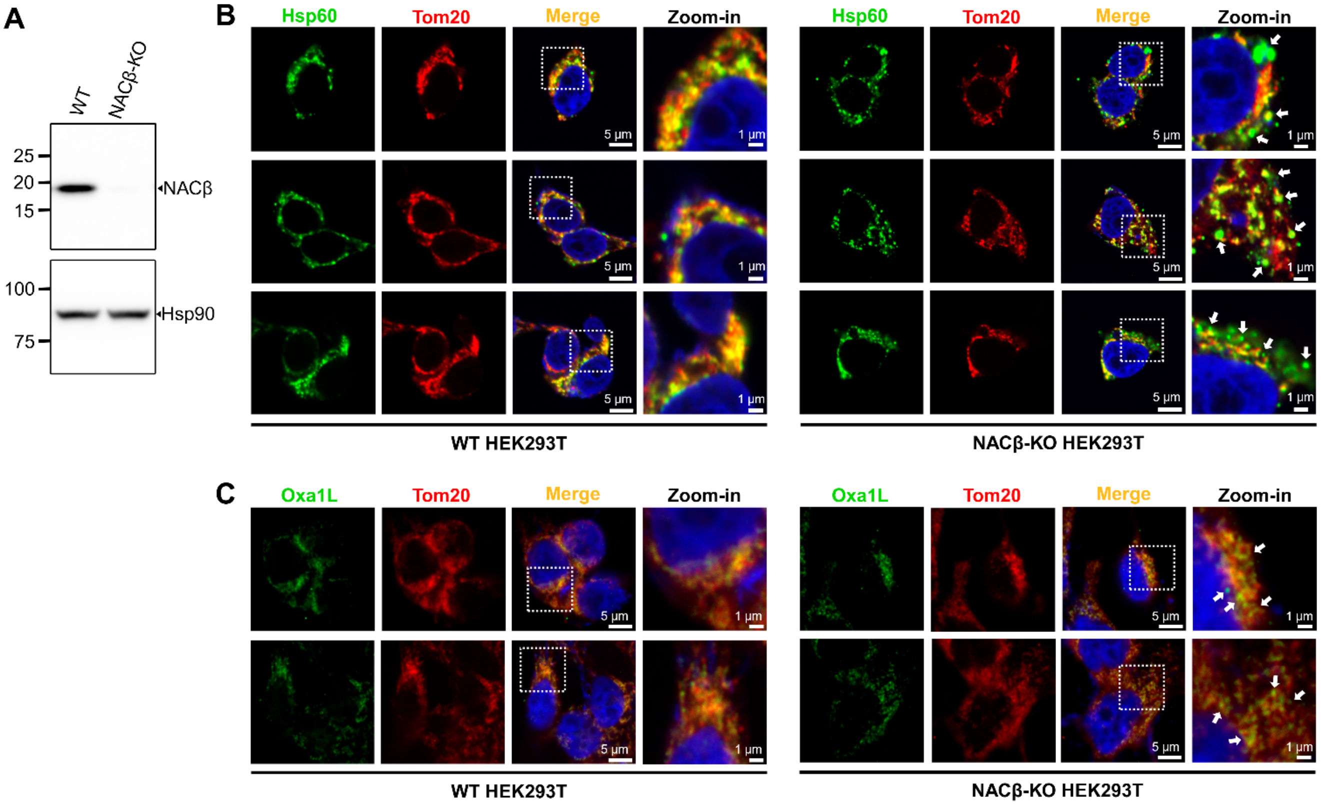


**Extended Data Figure 1.** Mitochondrial proteins are mislocalized in NACβ-KO HEK293T cells. A) WB depicting NACβ knocked out in HEK293T cells. Hsp90 is used as a loading control. B) Immunofluorescence staining for mitochondrial matrix chaperone Hsp60 in WT and NACβ-KO HEK293T cells. C) Immunofluorescence staining for inner mitochondrial membrane insertase Oxa1L in WT and NACβ-KO HEK293T cells. The outer mitochondrial membrane receptor Tom20 is used as a mitochondrial marker.

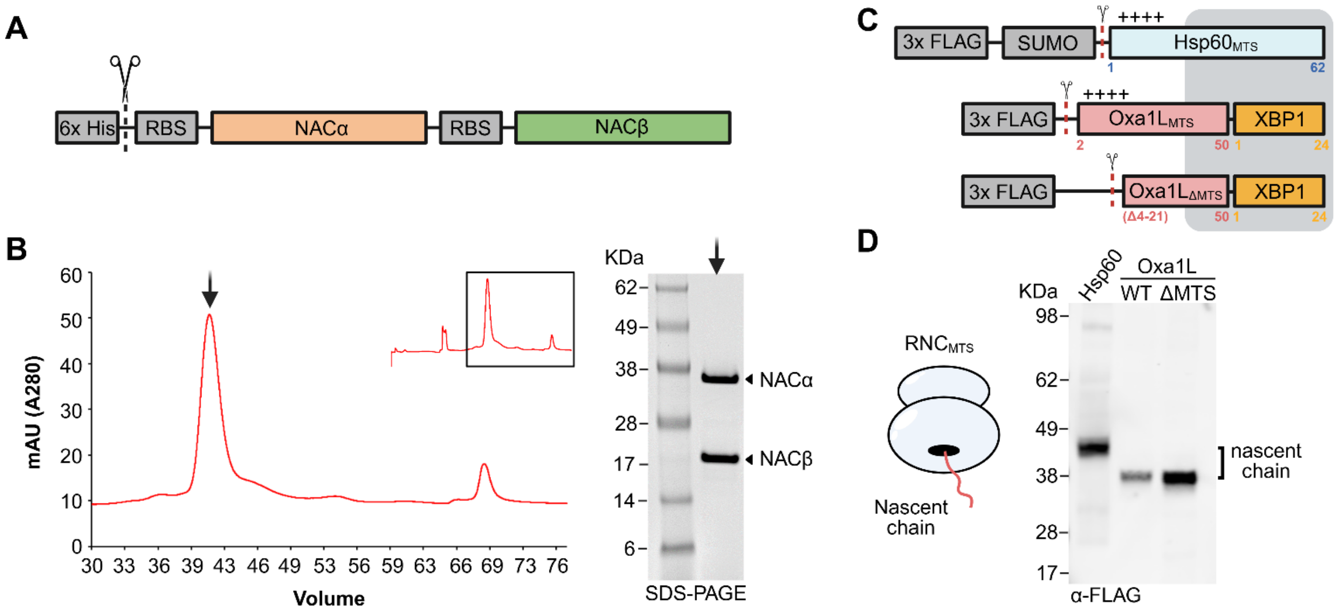


**Extended Data Figure 2.** Purification of the NAC complex and RNCs. **A)** Schematic of the construct used to recombinantly co-express both subunits of the NAC complex. **B)** Chromatogram from ion exchange purification of NAC indicating the peak fraction pooled and analyzed by SDS-PAGE. **C)** Schematic of the model mitochondrial nascent chains and MTS-deletion nascent chain purified for structural studies. **D)** Western blot validation of the purified translating ribosomes.

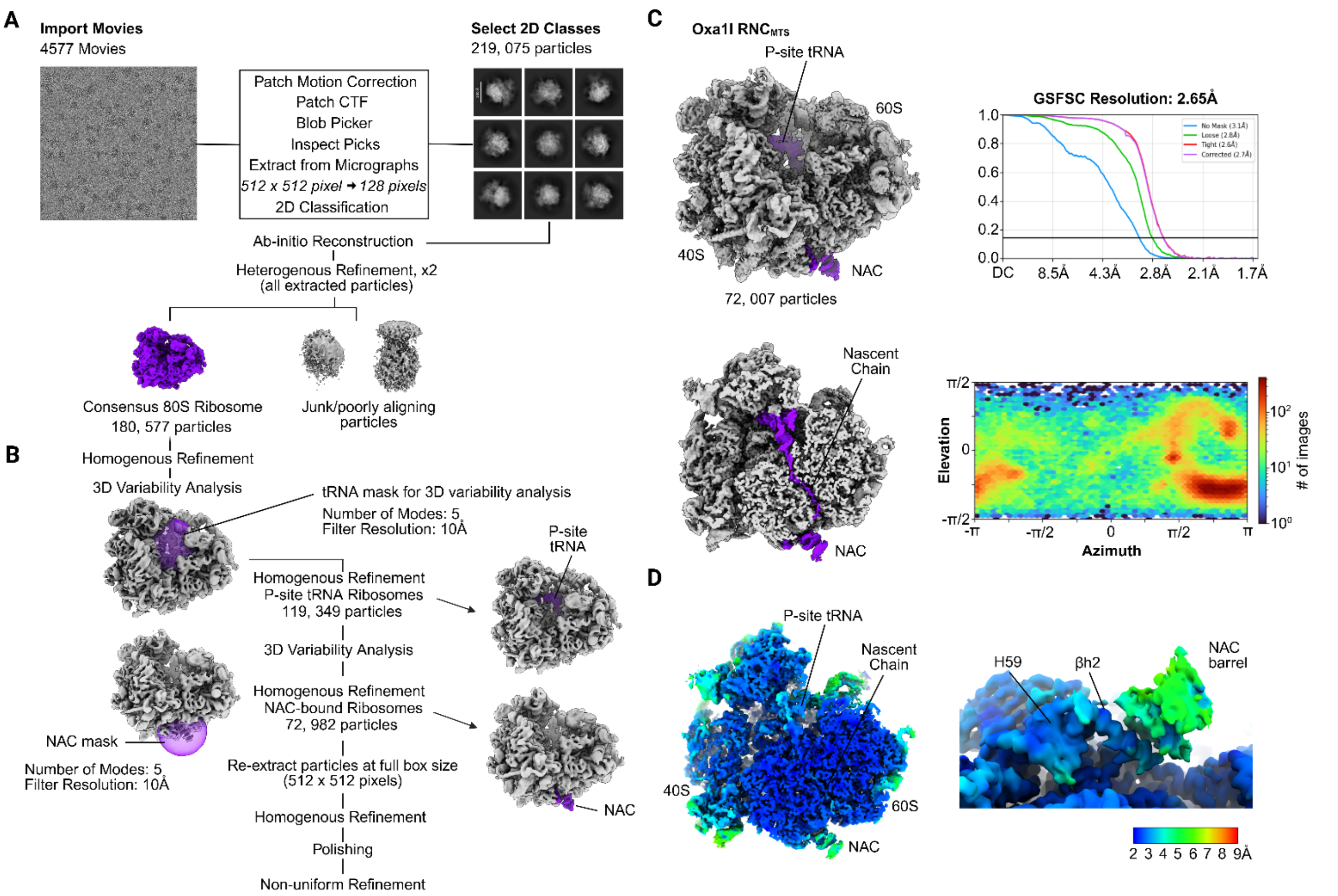


**Extended Data Figure 3.** Processing scheme for the structure of NAC in complex with ribosomes translating the N-terminal MTS of Oxa1L. **A)** Movies were motion and CTF corrected before particle picking and 2D classification. Ab-initio reconstruction of particles and subsequent heterogenous refinements were used to sort for 80S ribosomes and remove poorly aligning or junk particles form the dataset. **B)** Iterative 3D variability analyses on 80S ribosomes were conducted using focused spherical masks (purple) to sort for particles with strong densities for P-site tRNA and NAC. **C)** The particles with the best NAC and tRNA density were refined separately. The top right graph depicts the GSFSC curves between two independently reconstructed half-maps using an FSC cutoff of 0.143. Bottom right graph depicts the distribution of particle orientations in the dataset. **D)** Local resolution of the Oxa1L NAC-$RNC_{MTS}$ structure was determined using CryoSPARC at an FSC threshold of 0.143.

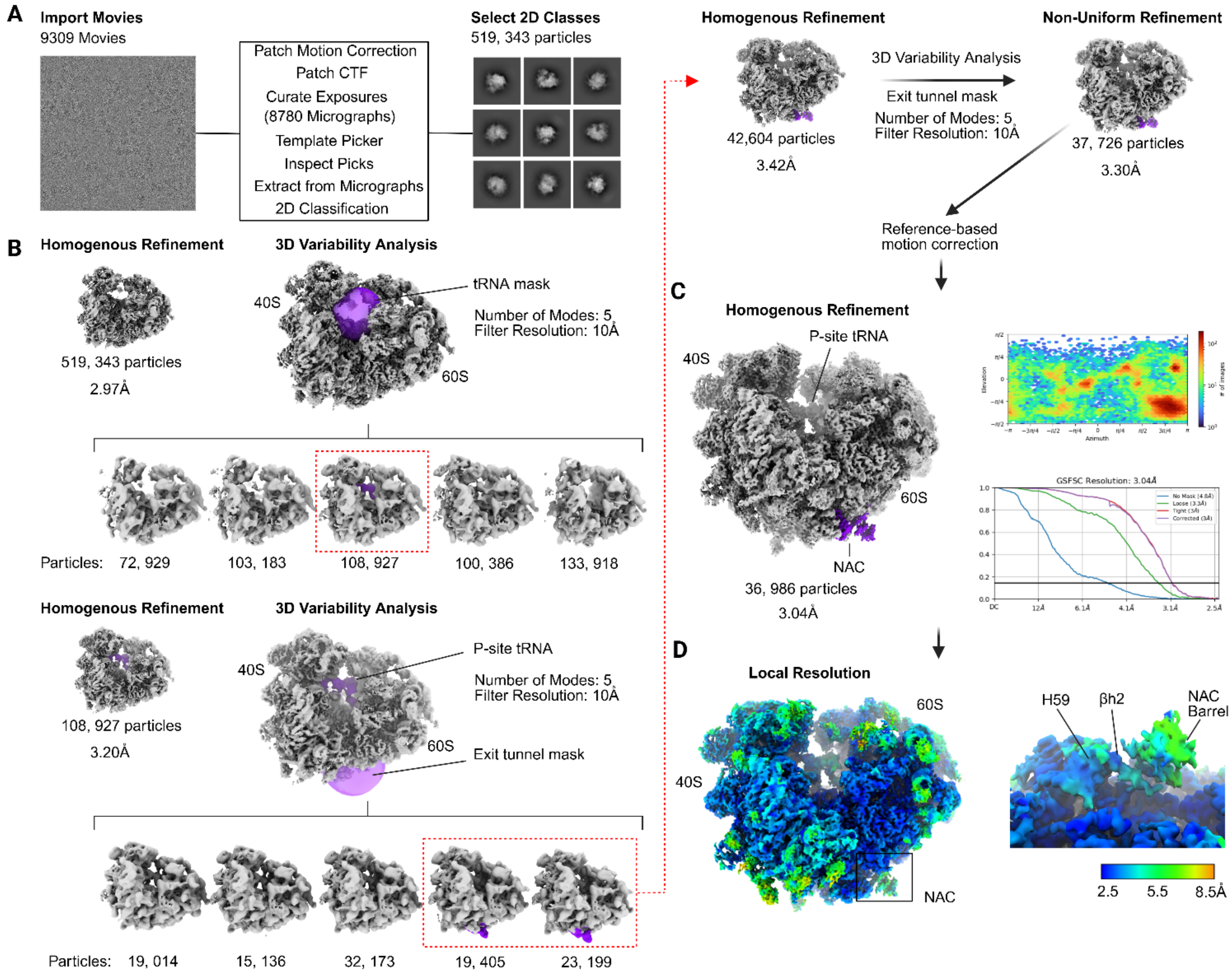


**Extended Data Figure 4.** Processing strategy and focused 3D classification scheme of NAC in complex with ribosomes translating the N-terminal MTS of Hsp60. **A)** Movies collected for single-particle analysis were motion and CTF corrected before particle picking and 2D classification. **B)** The particles selected following 2D classification were initially refined to generate a consensus structure of the 80S mammalian ribosome. Consensus 80S particles were further classified by conducting a series of sequential 3D variability analyses using focused spherical masks to sort for particles with strong NAC and P-site tRNA density. **C)** The final set of particles with the strongest NAC and tRNA density were refined separately. Top right graph shows the distribution of particle orientations in the dataset. Bottom right graph shows the gold standard Fourier shell correlation (GSFSC) determined between independently reconstructed half-maps using an FSC cutoff of 0.143. **D)** Local resolution of the complex was determined using CryoSPARC.

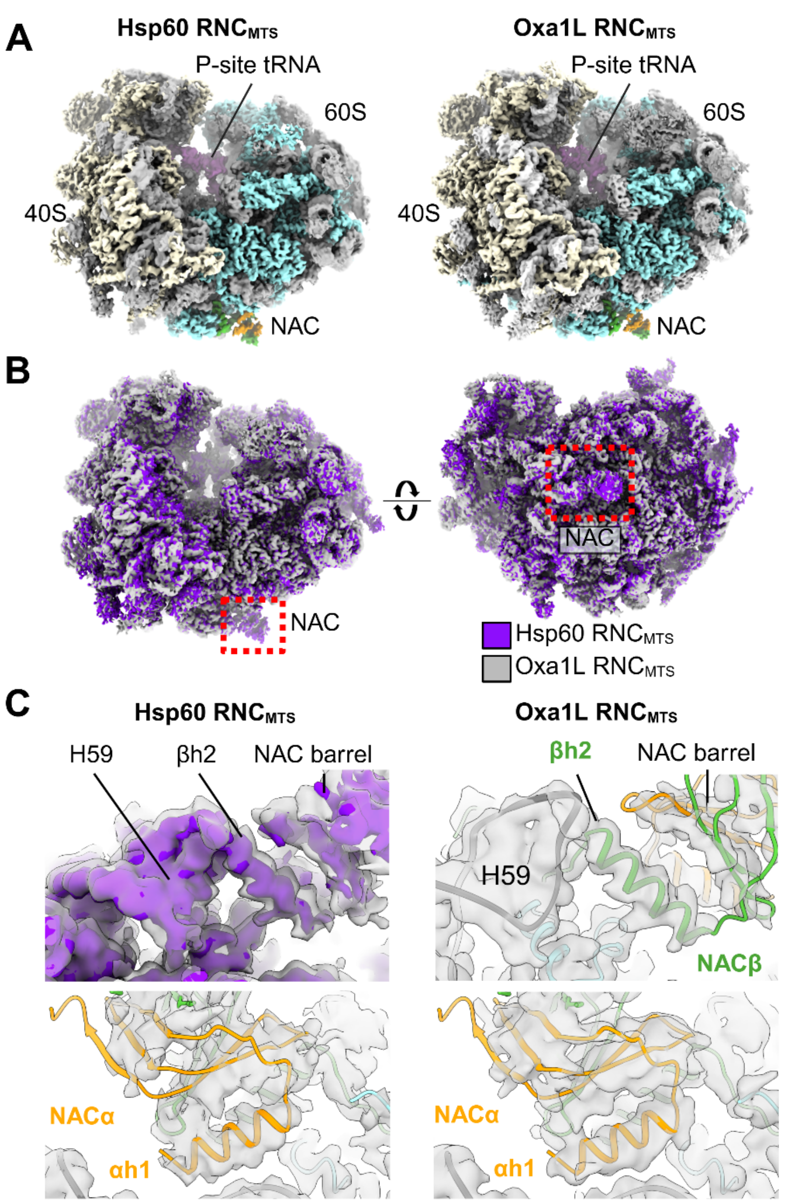


**Extended Data Figure 5.** Comparison of the Hsp60 and Oxa1L RNC$_{MTS}$ maps. **A)** Cryo-EM maps of RNCs carrying the N-terminal MTS of Hsp60 (left) and Oxa1L (right) filtered to 4 Å. **B)** Overlay of the Hsp60 (purple) and Oxa1L (grey) RNC$_{MTS}$ maps. **C)** Closeup of the overlay of the Oxa1L map with the Hsp60 map. The NAC-RNC$_{MTS}$ model was fit as a rigid body in the EM density for the indicated structure.

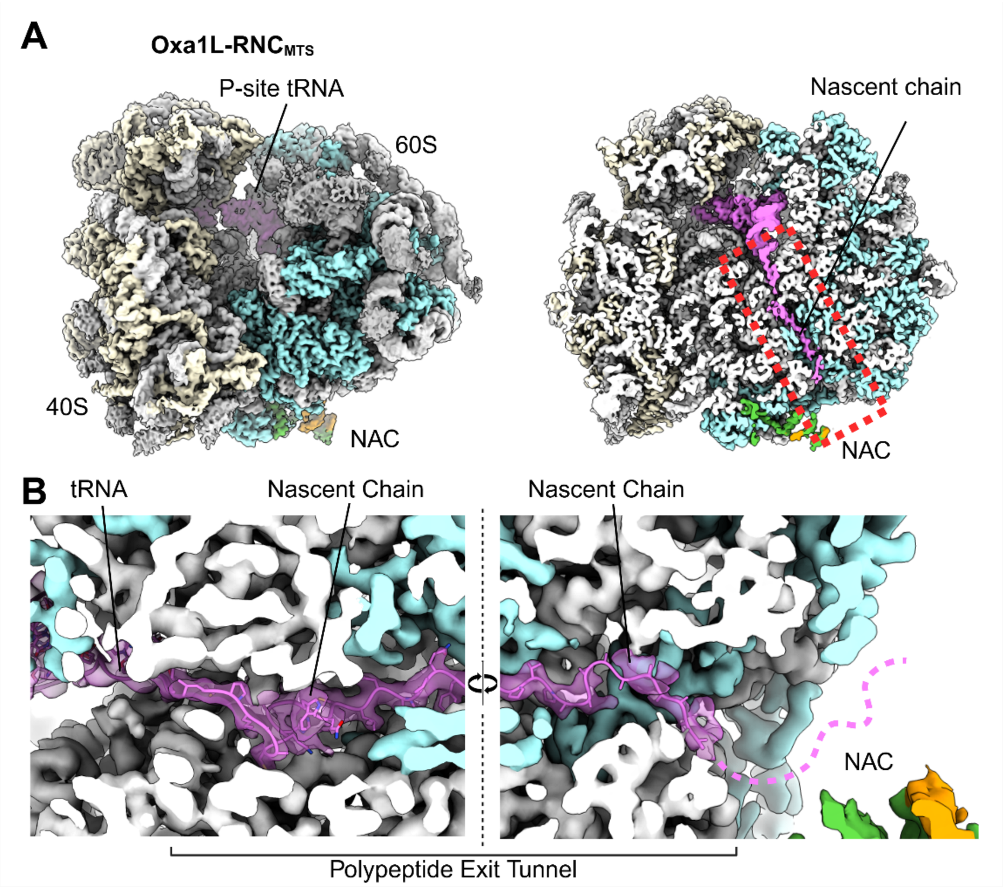


**Extended Data Figure 6.** Nascent chain density for the Oxa1L NAC-RNC$_{MTS}$ complex. **A)** An overview of the Oxa1L NAC-RNC$_{MTS}$ Cryo-EM map accompanied by a cross section showing the nascent chain within the polypeptide exit tunnel. **B)** Closeup of the nascent chain in the polypeptide exit tunnel for the boxed region in (A). The nascent chain density is shown as a transparent surface, with the underlying atomic model displaying resolved side chains of the stalled nascent chain. Colors: large ribosomal protein- cyan; small ribosomal proteins- beige; P-site tRNA and nascent chain- pink; NACα and NACβ- orange and green, respectively.

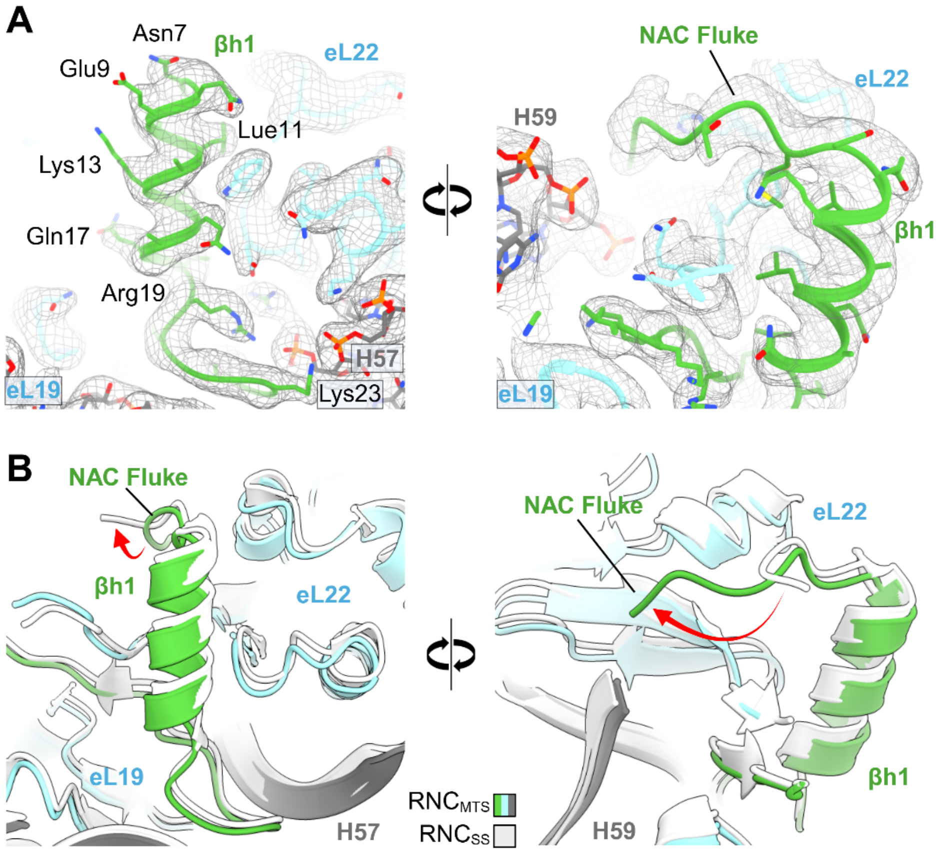


**Extended Data Figure 7.** Structure of the NACβ N-terminal fluke and anchor domain. **A)** Closeup of NAC βh1 (anchor) and N-terminus (fluke) depicting contacts with ribosomal proteins and rRNA. The cryo-EM map is shown as mesh. **B)** Comparison of the NACβ anchor domain and fluke between the NAC-RNC$_{MTS}$ structure and the NAC-RNC$_{SS}$ structure (PDB 7QWR). Red arrow indicates a conformational change in the fluke.

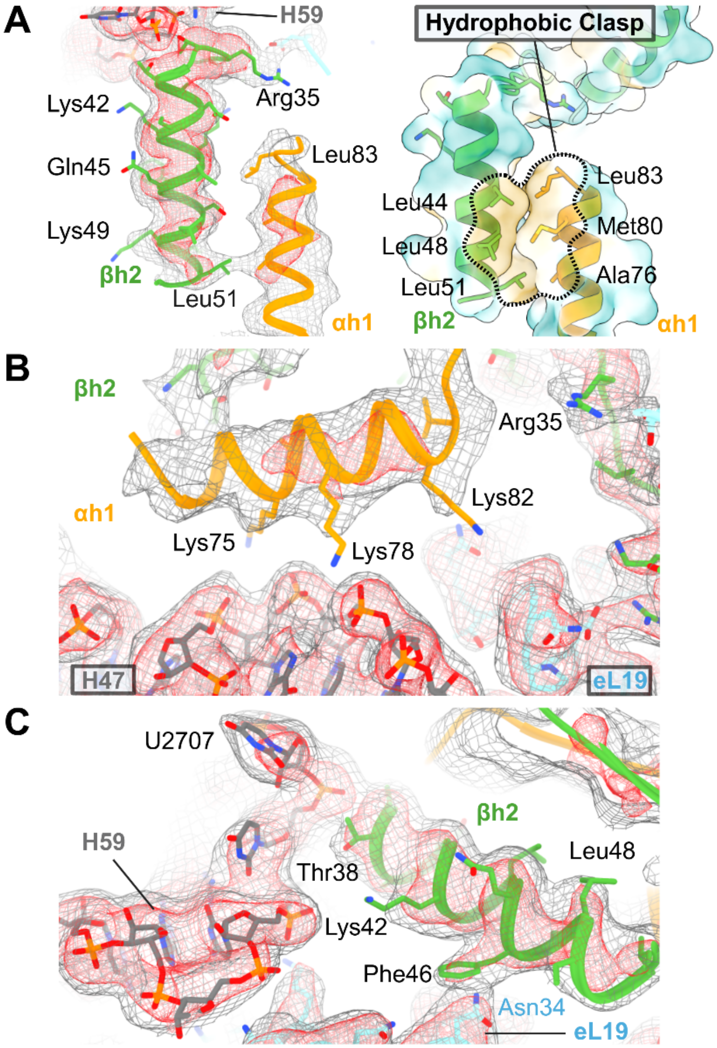


**Extended Data Figure 8.** Interactions of the NAC amphipathic helices with the ribosome. **A)** Map and model of the hydrophobic clasp (left) formed by the amphipathic helices of the NAC barrel domain and corresponding surface model colored by hydrophobicity (right). **B)** Closeup of NACα h1 interactions with rRNA H47. **C)** Side view of NAC βh2 interactions with H59 and eL19. The cryo-EM maps are depicted as mesh at 3.5 Å (red) and 4Å (grey) at two different thresholds for clarity.

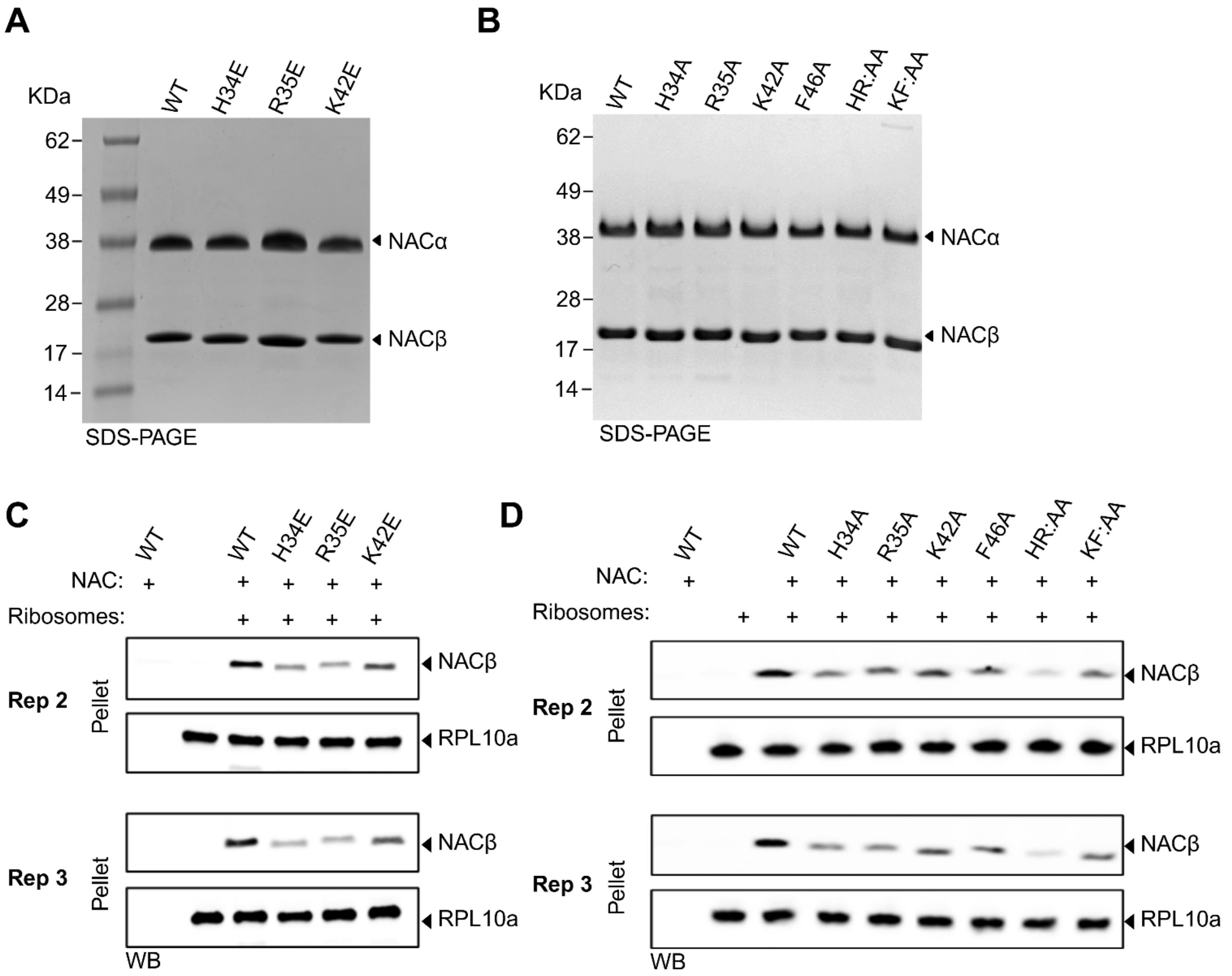


**Extended Data Figure 9.** NACβ switch mutants influence NAC binding to the ribosome. **A)** SDS-PAGE analysis of purified wild-type and indicated NAC glutamic acid (E) mutants. **B)** SDS-PAGE of purified wild-type and indicated NAC Ala (A) mutants. **C)** Additional blots used for the band quantification in Fig 2C of the co-sedimentation assays conducted with WT and glutamic acid NAC mutants. **D)** Additional blots used for the band quantification in Fig 2C-D co-sedimentation assay of NAC alanine variants. RPL10a was used as a loading control.

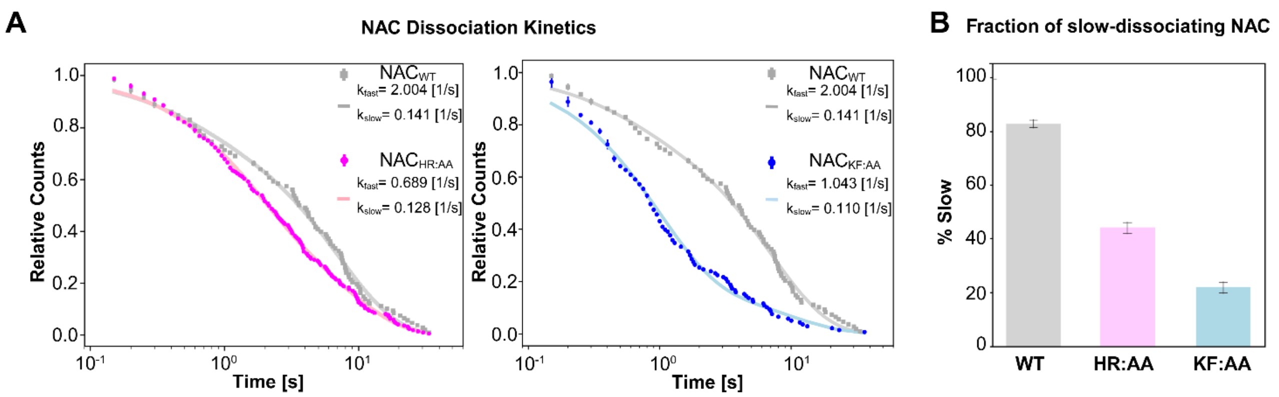


**Extended Data Figure 10.** NACβ switch mutants exhibit reduced kinetic stability on $RNC_{MTS}$. **A)** Kinetics of NAC dissociation from Oxa1L-$RNC_{MTS}$ is compared between WT NAC and $NAC_{HR:AA}$ (left) or $NAC_{KF:AA}$ (right). The lines show fits of the data to a double exponential function, which gave the indicated dissociation rate constants for the fast-dissociating ($k_{fast}$) and slow-dissociating ($k_{slow}$) populations. **B)** Summary of the fraction of slow-dissociating population for WT and mutant NAC, from fits of the data in A. Value are shown as fitted parameter ± standard error (SE) of the fit.

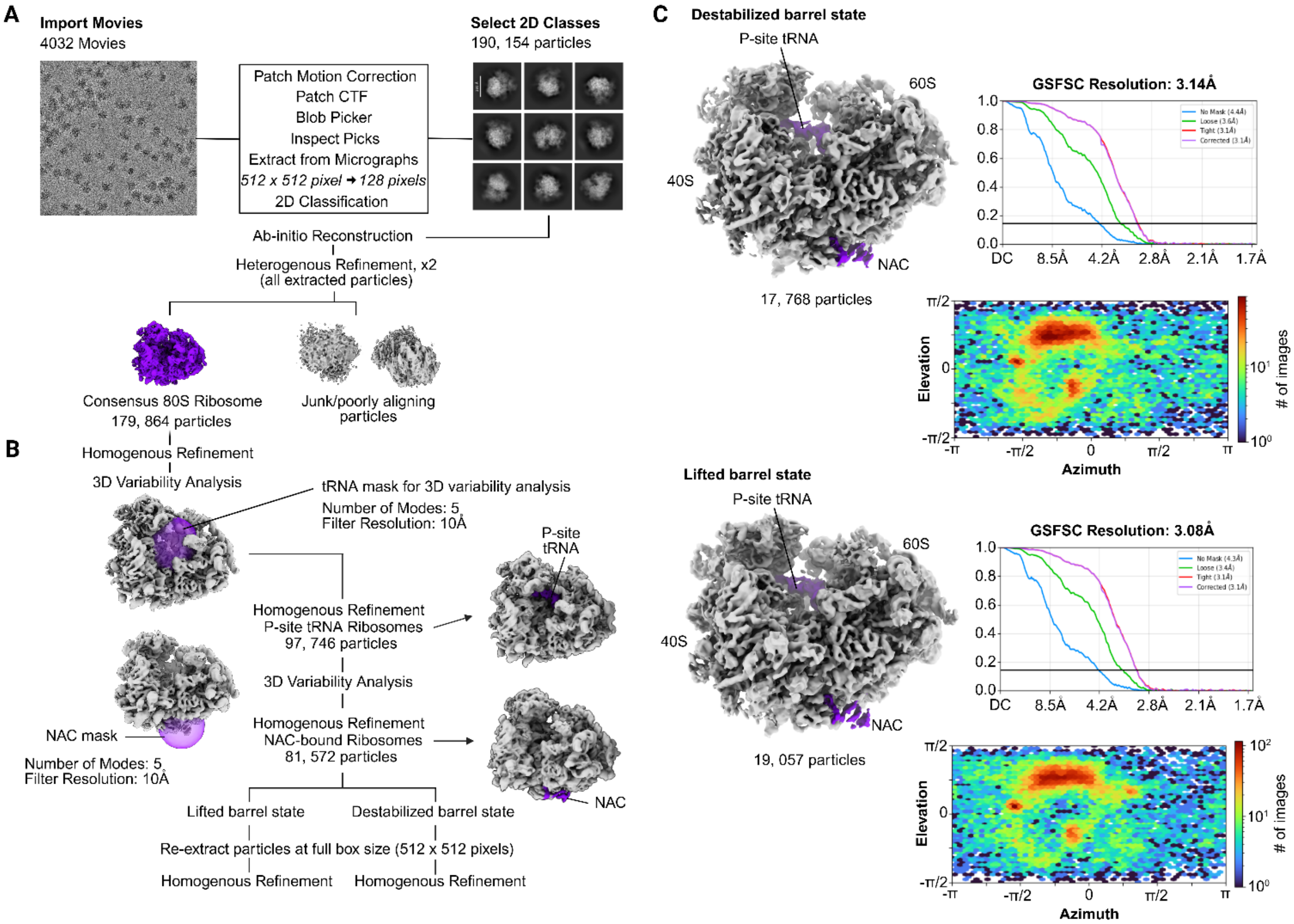


**Extended Data Figure 11.** Processing strategy and focused 3D classification scheme of NAC in complex with ribosomes translating the MTS deletion mutant of Oxa1L. **A)** Movies were motion and CTF corrected before particle picking and 2D classification. Selected particles underwent an initial 3D classification approach to remove junk and poorly aligning particles. **B)** A series of 3D variability analyses were conducted using focused spherical masks to sort for particles with P-site tRNA and NAC density. **C)** Particles corresponding to NAC barrel movements in a lowered (destabilized) and lifted (stabilized) conformation were refined separately. Top right graphs show the gold standard Fourier shell correlation (GSFSC) determined between independently reconstructed half-maps using an FSC cutoff of 0.143. Bottom right graphs show the distribution of particle orientations in the dataset.

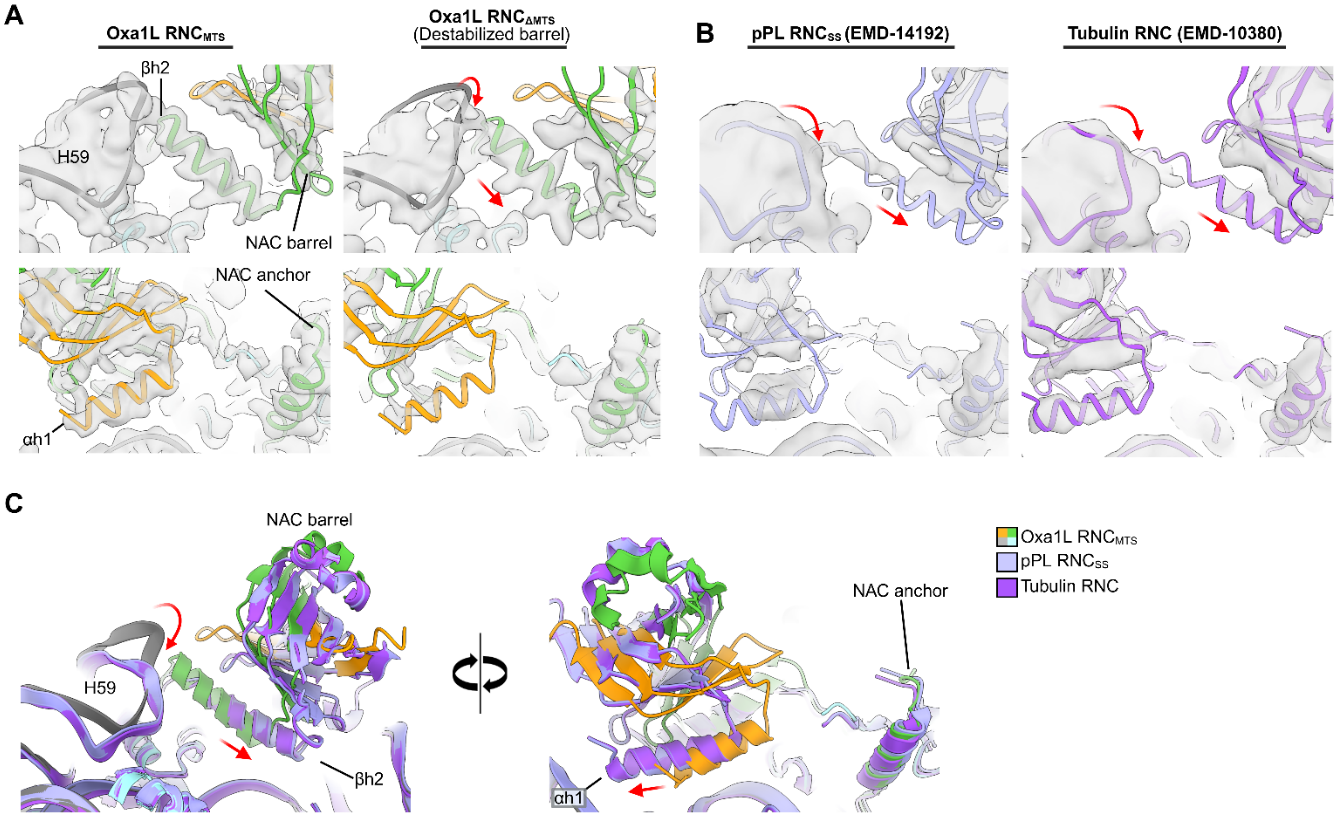


**Extended Data Figure 12.** Map and model comparison of NAC-RNC$_{MTS}$ to previous NAC structures on ribosomes translating SS$_{pPL}$ and tubulin. A) Fits of the NAC-RNC$_{MTS}$ model to the Oxa1L RNC$_{MTS}$ and Oxa1L RNC$_{\Delta MTS}$ maps. Closeups of the amphipathic helices of NAC (βh2 and αh1) are shown. B) Previous structures of NAC bound to RNCs translating SS$_{pPL}$ (PDB 7QWR) and tubulin (PDB 7QWS) fitted to corresponding maps, EMD-14192 and EMD-10380, respectively. Closeups of the amphipathic helices of NAC (βh2 and αh1) are shown with red arrows indicating conformational changes relative to the NAC-RNC$_{MTS}$ structure. C) Comparison of the conformational changes observed for the NAC barrel on ribosomes translating SS$_{pPL}$ (lavender) and tubulin (purple) nascent chains relative to ribosomes translating the Oxa1L MTS.

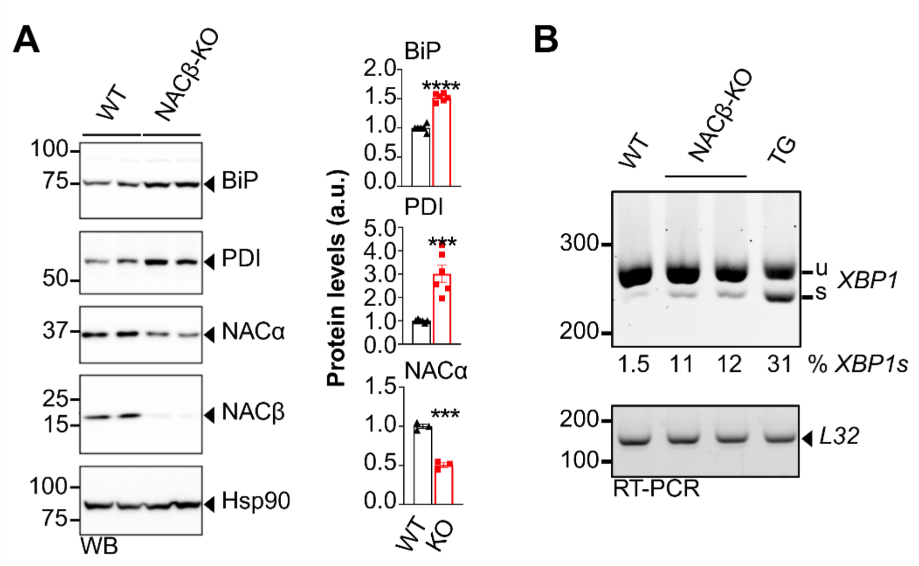


**Extended Data Figure 13.** NACβ knockout induces modest ER stress that elevates ER resident chaperones and increases the splicing of *XBP1* mRNA. **A)** Western blot for ER chaperones, BiP and PDI, following NACβ knockout. Bars show quantification of bands normalized to control cells. Hsp90, loading control. **B)** RT-PCR of *XBP1* splicing levels in WT and NACβ knockout HEK293T cells. The percentage of the ratio of spliced to total *XBP1* is shown below the gel. L32, loading control. TG, cells treated with Thapsigargin, a severe inducer of ER stress. An unpaired t-test was performed to assess statistical significance (***p < 0.001 and ****p< 0.0001).

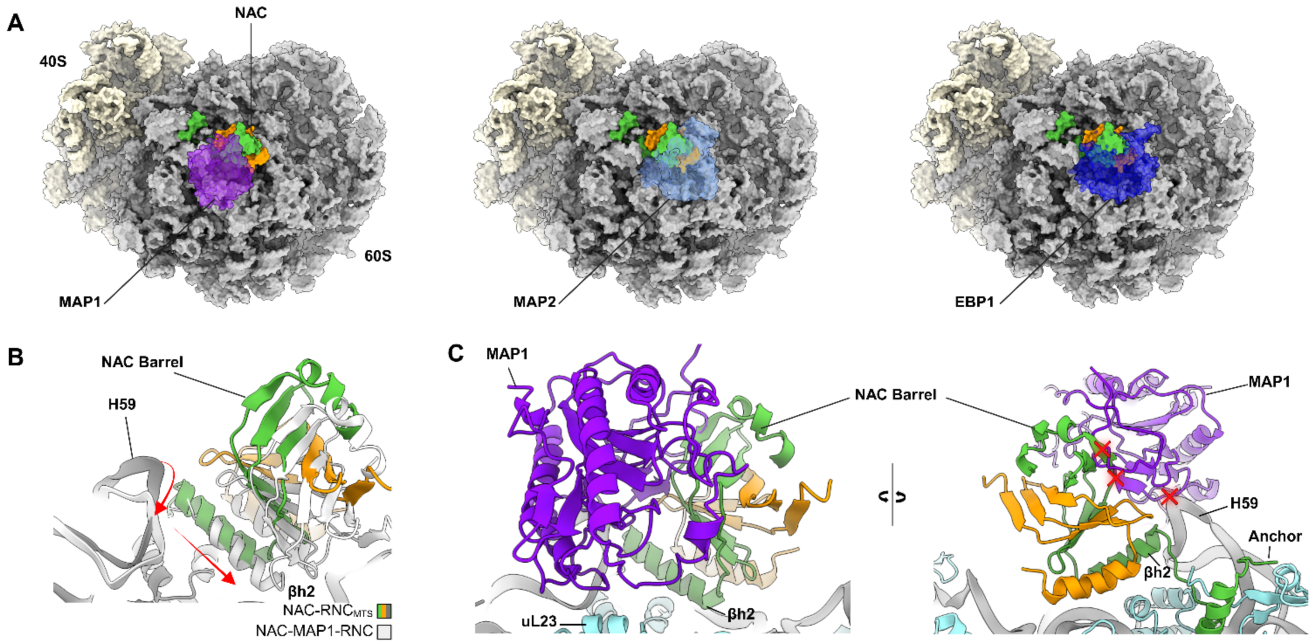


**Extended Data Figure 14.** Comparison of the NAC-RNC$_{MTS}$ structure with the previous MAP and EBP1 structures. **A)** Docking of MAP1 (PDB 8P2K), MAP2 (PDB 8ONY), and EBP1 (PDB 6SXO) to the NAC-RNC$_{MTS}$ structure. B) A closeup of the observed conformational change of the NAC barrel and H59 (red arrows) between the NAC-RNC$_{MTS}$ and NAC-MAP1-RNC structure is shown. **C)** Closeup of MAP1 and the NAC-RNC$_{MTS}$ structure highlighting clashes observed with a red 'X'.

**Extended Data Table 1.** Cryo-EM data collection, refinement and validation statistics.

| | Oxa1L $RNC_{MTS}$ | Hsp60 $RNC_{MTS}$ | Oxa1L $RNC\Delta_{MTS}$ | |
|---|---|---|---|---|
| | | | Destabilized barrel | Lifted barrel |
| EMDB code | EMD-48552 | EMD-71310 | EMD-71286 | EMD-71287 |
| PDB code | 9MR4 | | | |
| **Data collection and processing** | | | | |
| Nominal magnification | 105,000x | 120,000x | 105,000x | 105,000x |
| Voltage (kV) | 300 | 200 | 300 | 300 |
| Electron exposure ($e^{-}/Å^2$) | 32 | 50 | 40 | 40 |
| Defocus range (µm) | -1.4/-0.8 | -2.2/-1.2 | -1.4/-0.8 | -1.4/-0.8 |
| Pixel size (Å) | 0.833 | 1.17 | 0.83 | 0.83 |
| Initial particle images (no.) | 180,577 | 519,343 | 179,864 | 179,864 |
| Final particle images (no.) | 72,007 | 36,986 | 17,768 | 19,057 |
| Map resolution at FSC=0.143 | 2.65 | 3.04 | 3.14 | 3.08 |
| **Structure refinement in PHENIX 1.20.1** | | | | |
| Model resolution at FSC=0.5 | 2.9 | | | |
| $CC_{mask}$ | 0.82 | | | |
| Map sharpening B factor ($Å^2$) | 60.8 | | | |
| Model composition | | | | |
| Non-hydrogen atoms | 211,175 | | | |
| Protein residues | 11,534 | | | |
| RNA residues | 5,513 | | | |
| B factors min/max/mean ($Å^2$) | | | | |
| Protein | 0.00/53.84/24.07 | | | |
| RNA | 0.00/82.57/28.67 | | | |
| Ligand | 0.00/69.18/19.44 | | | |
| RMSD | | | | |
| Bond lengths (Å) | 0.003 | | | |
| Bond angles (°) | 0.719 | | | |
| **Validation** | | | | |
| MolProbity score | 1.73 | | | |
| Clashscore | 7.46 | | | |
| Poor rotamers (%) | 1.98 | | | |
| Ramachandran plot | | | | |
| Favored (%) | 97.55 | | | |
| Allowed (%) | 2.43 | | | |
| Outliers (%) | 0.02 | | | |
